# Neuron-radial glial cell communication via BMP/Id1 signaling maintains the regenerative capacity of the adult zebrafish telencephalon

**DOI:** 10.1101/2021.05.26.445748

**Authors:** Gaoqun Zhang, Luisa Lübke, Fushun Chen, Tanja Beil, Masanari Takamiya, Nicolas Diotel, Uwe Strähle, Sepand Rastegar

## Abstract

The mechanisms of the management of the neural stem cell (NSC) pool underlying the regenerative capacity of the adult zebrafish brain are not understood. We show that Bone Morphogenetic Proteins (BMPs) which are exclusively expressed by neurons in the adult telencephalon and the helix-loop-helix (HLH) transcription co-regulator, Inhibitor of differentiation 1 (Id1), control quiescence of NSCs. Upon injury, lack of *id1* function leads to an initial over-proliferation and subsequent loss of NSCs and the regenerative capacity. BMP/Id1 signaling up-regulates the transcription factor *her4.1* which is also a target of Notch signaling mediating short-range control of NSC quiescence. Hence, the two signaling systems converge onto Her4.1. Our data show that neurons feedback on NSC proliferation. BMP1/Id1 signaling appears as the predominant safeguard of the NSC pool under regenerative conditions while Notch signaling is sufficient to maintain NSCs under homeostatic baseline neurogenesis in the uninjured animal.

## Introduction

Adult mammals have a limited ability to generate new neurons and to repair injured nervous tissue (Alunni and Bally-Cuif, 2016). In contrast, the brain of adult zebrafish contains many distinct neurogenic niches (Adolf et al., 2006; Grandel et al., 2006; Lindsey et al., 2018). In addition, and also unlike mammals, zebrafish can repair large neural lesions efficiently, frequently recovering function without striking disabilities (Adolf et al., 2006; Becker and Becker, 2008; Kroehne et al., 2011; März et al., 2011; Zupanc and Sirbulescu, 2011). In the telencephalon of adult zebrafish, the entire ventricular zone produces new neurons under physiological conditions representing baseline or constitutive neurogenesis (Adolf et al., 2006; Diotel et al., 2020; Ghaddar et al., 2021; Kroehne et al., 2011; März et al., 2011; Pellegrini et al., 2007). When the telencephalon is injured, proliferation of NSCs is transiently boosted above the baseline of constitutive neurogenesis with a peak at five to seven days after injury (Diotel et al., 2013; Ghaddar et al., 2021; März et al., 2011; Simon et al., 2011). Peculiarly, injury inflicted within one hemisphere of the telencephalon leads to a proliferative response of NSCs only in this injured hemisphere while no response is observed in the closely apposed stem cell niche of the uninjured half of the telencephalon (März et al., 2011; Rodriguez Viales et al., 2015; Zhang et al., 2020). Thus, the signals which trigger stem cell proliferation in response to injury remain confined within the injured hemisphere.

The ventricular zone of the adult zebrafish telencephalon is densely populated by the cell bodies of radial glia cells (RGCs), - the NSCs of the zebrafish telencephalon (März et al., 2010; Pellegrini et al., 2007). RGCs express typical NSC-markers such as Glial acidic fibrillary protein (Gfap), Brain lipid binding protein (Blbp), and the Calcium-binding protein β (S100β) (Diotel et al., 2020; Diotel et al., 2016; Ghaddar et al., 2021; Lam et al., 2009; März et al., 2010). Under homeostatic conditions, the majority of the NSCs are quiescent type I RGCs. Only a small proportion of NSCs proliferate and express proliferation markers, such as proliferating cell nuclear antigen (PCNA). This latter so-called type II RGCs can give rise to committed neuronal progenitors corresponding to neuroblasts (type III cells) (Ghaddar et al., 2021; März et al., 2010). When the telencephalon is injured, many more RGCs enter the cell cycle and start to express proliferation markers (März et al., 2011). Concomitantly, the RGCs generate an increased number of new neurons compared to homeostatic conditions (Barbosa et al., 2015). Newborn neuronal precursors migrate from the ventricular layer to the injury site to replace lost neurons (Kroehne et al., 2011). This regenerative neurogenesis can be initiated by inflammatory signals (Kizil et al., 2012; Kyritsis et al., 2014; Kyritsis et al., 2012).

During regeneration, NSCs/RGCs were proposed to divide mostly symmetrically giving rise to two new neurons or two RGCs (Barbosa et al., 2015). Especially, the shift to regenerative neurogenesis in response to injury would rapidly lead to a depletion of stem cells unless these fate decisions are carefully managed. Genetical and pharmacological evidence strongly support a role of Notch signaling in regulating NSC quiescence, both in mouse and zebrafish (Chapouton et al., 2010; Ehm et al., 2010; Imayoshi et al., 2010; Kawai et al., 2017). Specifically, in the zebrafish telencephalon Notch3 is expressed in RGCs and promotes quiescence of NSCs (Alunni et al., 2013). This signaling originates, at least in part, from the progenitor population itself (Dray et al., 2021a; Dray et al., 2021b). Notch signaling appears to serve as an intrinsic signaling cue for the regulation of quiescence and the maintenance of the stem cell population in mammals and fish (Diotel et al., 2020; Alunni and Bally-Cuif, 2016).

Previously, in a genome-wide search for transcription regulator genes differentially expressed in response to telencephalic injury in the zebrafish (Diotel et al., 2015a; Diotel et al., 2015b; Rodriguez Viales et al., 2015), we identified the HLH factor Id1 which is a non-DNA-binding inhibitor of basic-Helix-Loop-Helix (bHLH) transcription factors (Roschger and Cabrele, 2017). *id1* was shown to be mainly expressed in quiescent RGCs and to be specifically up-regulated in these cells after injury (Rodriguez Viales et al., 2015; Zhang et al., 2020). As in constitutive neurogenic condition, *id1* expression was mainly associated with non-proliferative RGCs after brain injury (Rodriguez Viales et al., 2015). Functional studies using morpholinos and mosaic gain-of-function by *in situ* lipofection suggested a role of Id1 in repressing proliferation of NSCs. We speculated that in particular its up-regulation after injury is important to maintain the NSC pool after injury (Rodriguez Viales et al., 2015).

Expression of *id1* in RGCs is driven by a phylogenetically conserved *cis*-regulatory module (CRM) (Zhang et al., 2020). For activity under both constitutive and regenerative conditions, this CRM requires Smad1/5 and Smad4 binding motifs which are intracellular mediators of BMP signaling (Zhang et al., 2020). Transcriptome analysis, as well as pharmacological inhibition experiments supported a possible role of BMP signaling for the function of the CRM. Our preliminary data therefore suggested that, in addition to Notch, BMP signaling may play a role in regulating the proliferative activity of NSCs during constitutive and regenerative neurogenesis in the zebrafish telencephalon (Zhang et al., 2020).

Here, we tested the hypotheses (1) that BMP signaling represses proliferation of NSCs in the zebrafish telencephalon and (2) that this repression is required to maintain stem cell pools and thus the regenerative capacity of the adult zebrafish telencephalon. By conditional genetic approaches, we show that BMP signaling controls proliferation as well as *id1* expression, consistent with the notion that BMPs/Id1 are regulators of stem cell quiescence. The BMP system operates independently and in parallel to Notch signaling and appears to converge on the expression of the Notch downstream mediator *her4.1*. Moreover, genetic ablation of *id1* function leads to an increased proliferation of NSCs and increased depletion of stem cell pools in injured mutant telencephala with concomitant failure to heal injuries. Our data show that BMP signaling is key to the maintenance of NSC pools and the long-term capacity to regenerate wounds in the zebrafish telencephalon. Moreover, our data suggest that in addition to the proposed short-range Notch signaling which is intrinsic to the stem cell niche, external cues from the regenerating tissue maintain the proliferation-competent stem cell pool in the zebrafish telencephalon.

## Results

### BMP proteins are expressed in neurons of the telencephalon

Preliminary data suggested that BMPs control expression of *id1* and in this way regulate cell proliferation in the telencephalon (Zhang et al., 2020). We first addressed the question of whether *bmps* are expressed in the telencephalon of the adult zebrafish. We chose *bmp2a*, *bmp2b*, *bmp4*, *bmp7a* and *bmp7b* for their key roles during early development of the zebrafish embryo (Kondo, 2007). We carried out expression analyses on transverse sections through the telencephalon by *in situ* hybridization (ISH) with either chromogenic (Figures 1A-E) or fluorescent staining (Figures 1F-J) of antisense probes directed against mRNAs of *bmp* genes. Expression of the mRNAs of the five *bmp* genes was detected in the brain parenchyma and along the telencephalic periventricular layer in similar and overlapping patterns, although levels of expression varied between different sub-regions of the telencephalon for individual *bmp* genes (Figures 1A-E).

**Figure 1.**
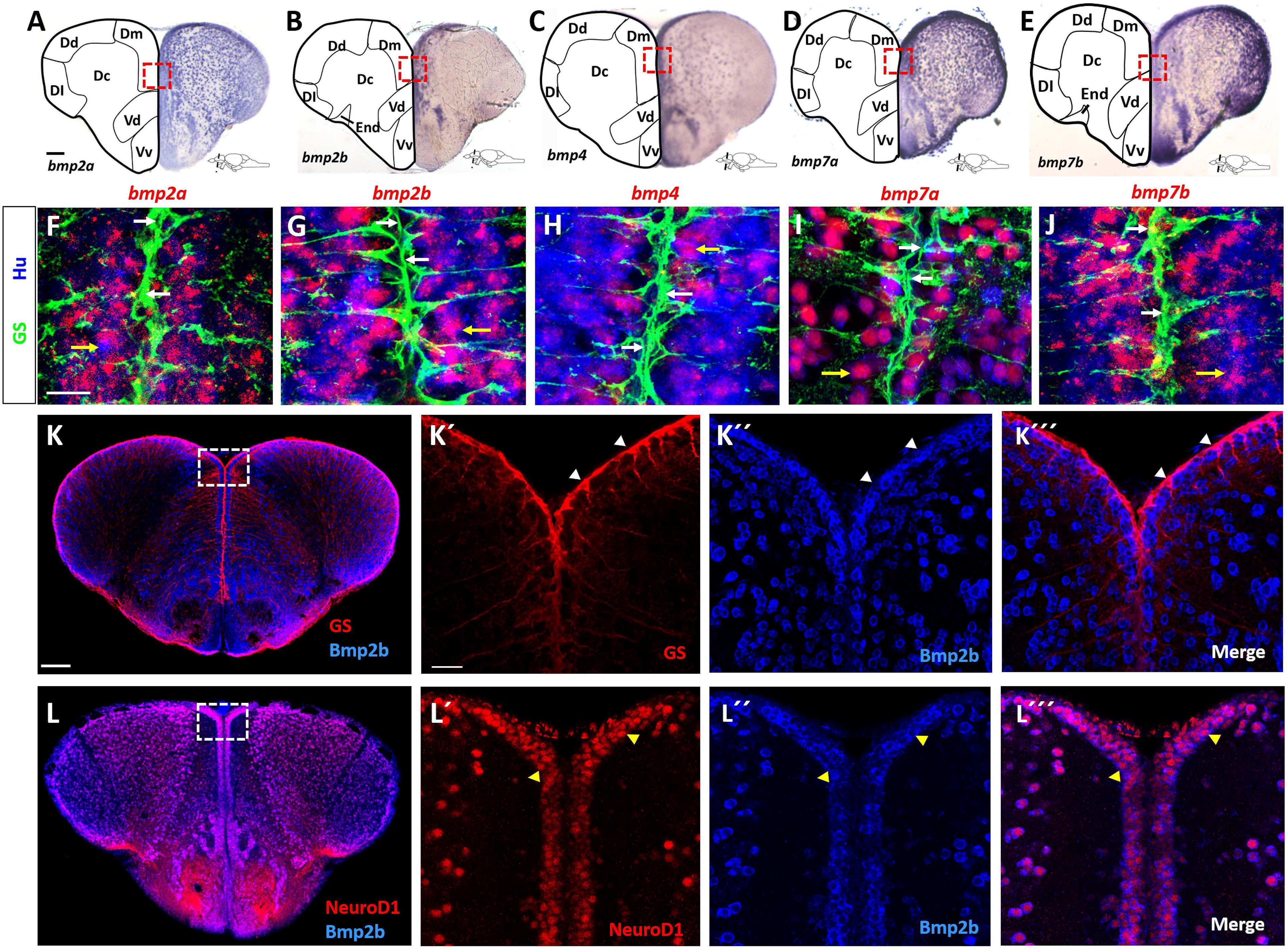
Expression of *bmp* genes in the adult zebrafish telencephalon. (A-E) *in situ* hybridization (ISH) with probes directed against *bmp2a* (A), *bmp2b* (B), *bmp4* (C), *bmp7a* (D) and *bmp7b* (E) mRNA on transverse telencephalic sections. All *bmp* genes show wide-spread expression. While *bmp2a*, *bmp7a* and *bmp7b* appear to be expressed in all telencephalic nuclei, *bmp2b* and *bmp4* mRNAs are more confined to the medial region. Red rectangles (A-E) indicate the coordinates of the region presented in F-J, respectively. (F-J) Expression of *bmp* genes revealed by fluorescent ISH (red) on cross-sections of WT telencephala, together with double immunofluorescent staining for RGCs (glutamine synthetase [GS], green) and post-mitotic neurons (Hu, blue). The *bmp* genes are co-expressed with Hu in neurons (yellow arrows in F-J) but are not expressed in GS+ RGCs (white arrows). (K-L’’’) BMP2b antibody staining (blue) with GS (K, red) or NeuroD1 (L, red) antibodies shows that Bmp2b is expressed in NeuroD1+ neurons. White rectangles (K, L) represent the region magnified in K’-K’’’, L’-L’’’, respectively. White arrowheads show the GS+ RGCs (K’-K’’’), yellow arrowheads indicate neurons (L’-L’’’). Scale bar = 20 μm (F-J, K’-K’’’, L’-L’’’), 100 μm (A-E, K, L). Dc: central zone of the dorsal telencephalic area; Dl: lateral zone of the dorsal telencephalic area; Dm: medial zone of the dorsal telencephalic area; Vc: central nucleus of the ventral telencephalic area; Vd: dorsal nucleus of the ventral telencephalic nucleus; Vv: ventral nucleus of the ventral telencephalic area.

Next, we tested whether neurons and/or RGCs express the *bmp* genes. To this end, we combined fluorescent *in situ* hybridization (FISH) for the five *bmp* probes with immunohistochemistry (IHC) using anti-HuC/D antibodies (Hu) to label post-mitotic neurons. By additional co-staining with antibodies directed against glutamin synthetase (GS), we marked RGCs in the telencephalic tissue sections. All five *bmp* genes were strongly expressed in neurons while their expression was not detectable in RGCs (Figures 1F-J). Thus, RGCs, with their cell bodies at the medial periventricular zones and their long processes traversing the parenchyma all the way to the pial surface, are embedded in a *bmp*-expressing, neuronal environment.

We next assessed whether this abundant expression of *bmp* mRNA in neurons also reflects the expression of the BMP protein. To address this, we conducted IHC using a BMP2b antibody and an antibody directed against NeuroD1. To assess expression in RGCs, we carried out double IHC with the BMP2b antibody and the GS antibody marking RGCs. BMP2b was not expressed in RGCs (Figures 1K-K’’’, white arrowheads). However, and as expected from the ISH data, BMP2b expression was associated with the NeuroD1 expressing neurons (Figures 1L-L’’’, yellow arrowheads). We also checked whether oligodendrocytes marked by *Tg(olig2:gfp)* (Park et al., 2007), co-express BMP2b. Only weak or absent BMP2b staining was detected in oligodendrocytes (Figure S1).

These data are consistent with a model where neurons are BMP-producing and RGCs are BMP-sensing cells in the telencephalon. To assess whether RGCs are, in principle, able to respond to BMP signals, we mined single cell sequencing data (Lange et al., 2020) for the expression of components of the BMP signaling pathway (Table S1). Quiescent RGCs were unequivocally identified by expression of *id1* and a lack of *cyclin D1* (*ccnd1*) transcripts (Lange et al., 2020). All 22 cells identified with this expression pattern expressed the BMP receptors *bmpr1ab* and *bmpr2b* and the downstream BMP specific mediators *smad1*, *smad5* and *smad9* (Table S1). Consequently, RGCs harbor the relevant components of the canonical BMP signaling pathway suggesting that they can respond to BMP signals. In addition, the BMP induced gene *bambia* is expressed in RGCs (Table S1) in agreement with the notion that RGCs receive BMP signals.

### Bmp signaling induces *id1* expression and promotes stem cell quiescence

To test the relevance of the observed BMP expression, we manipulated BMP signaling by conditionally inducing BMP signaling in adult zebrafish. We triggered Bmp2b expression with the help of a stably integrated, heat-shock inducible transgene *Tg(hsp70:bmp2b)* (Chocron et al., 2007). We checked the efficiency of the manipulation by employing an antibody directed against the phosphorylated states of Smad1, 5 and 9, the known intracellular mediators of the canonical BMP signaling cascade. Phosphorylation of these Smads reflects the activation of the serine/threonine kinase domains of the BMP receptors upon BMP ligand binding (Hata and Chen, 2016). No phosphorylated Smad1/5/9 (pSmad) staining was detectable above background in GS-marked RGCs in the absence of heat-shock (Figures 2B-B’’). In striking contrast, when BMP2b was ectopically expressed by heat-shock, strong immunoreactivity of pSmads was noted in RGCs (Figures 2C-C’’). Wild-type (WT) animals not carrying the *Tg(hs:bmp2b)* transgene did not show pSmad staining upon heat-shock (Figures 2A-A’’). Thus, ectopic expression of Bmp2b leads to a strong activation of BMP signaling in RGCs. Of note, the induction of phosphorylation of Smad is restricted to the RGCs, suggesting that neurons, the source of endogenous BMP signals, do not respond to BMPs in a strong fashion under the employed induction scheme. This further supports our model that neurons are BMP sending and RGCs are BMP perceiving cells.

**Figure 2.**
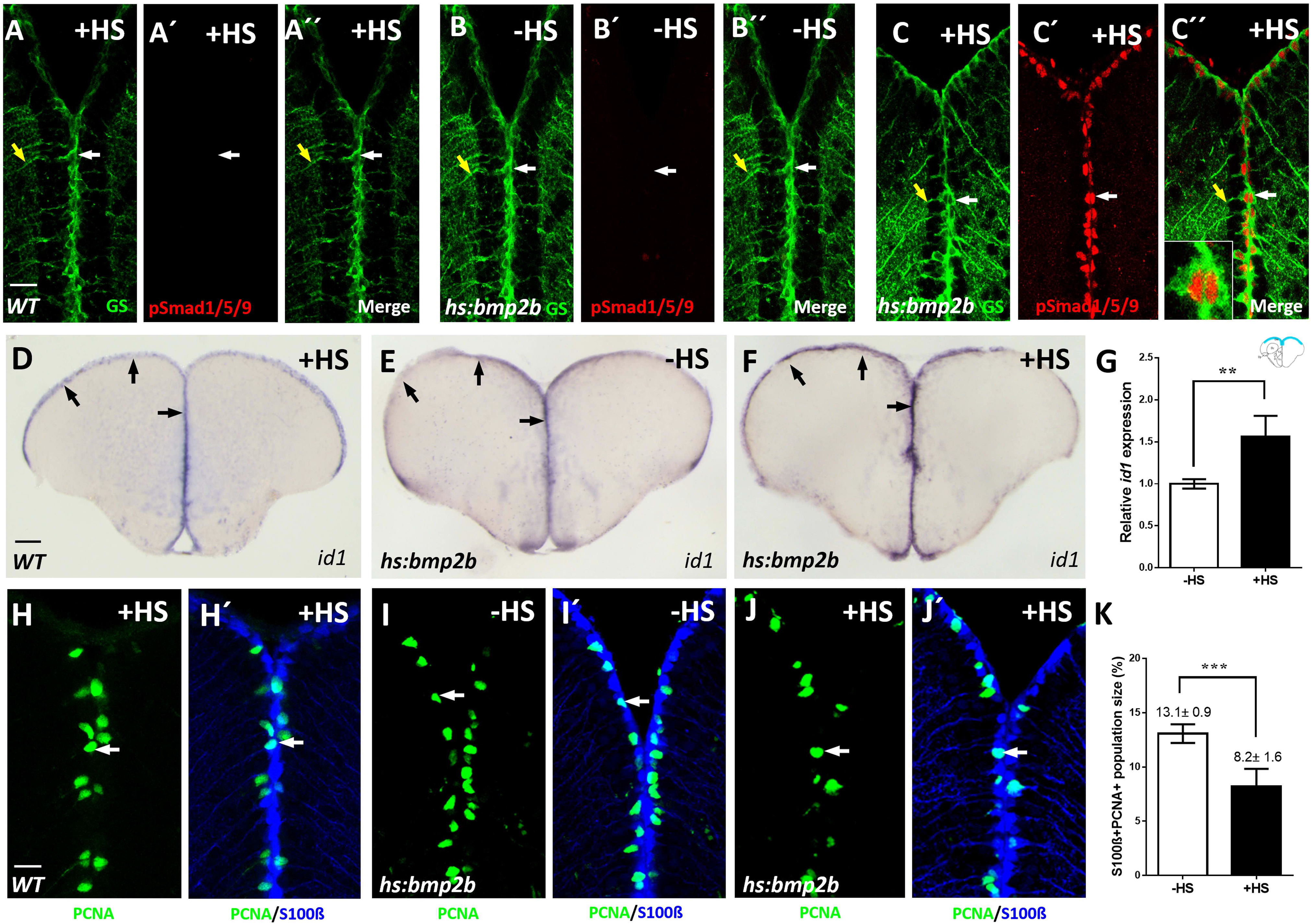
Conditional expression of *bmp2b* causes activation of BMP signaling, elevated expression of *id1* and an increased proliferation of RGCs. (A-C’’) Immunostaining with GS (green) and pSmad1/5/9 (red) antibodies on sections of WT telencephala with heat-shock (A-A’’, +HS), *Tg(hs:bmp2b)* telencephala without heat-shock (B-B’’, -HS) or with heat-shock (C-C’’, +HS). pSmad1/5/9 immunoreactivity was detected in GS+ RGCs exclusively in heat-shocked telencephala (n=3 brains). C’’ inset: a magnified view of a pSmad+/GS+ RGC. Note the nuclear localization of pSmad in contrast to cytoplasmic GS localization. White arrows show individual RGCs, yellow arrows indicate the processes of the same RGCs (A-C’’). (D-F) ISH showing increased *id1* expression after heat-shock-induced bmp2b expression. Black arrows show the expression of *id1* in the ventricular zone of WT (D), *Tg(hs:bmp2b)* after heat-shock (F) and *Tg(hs:bmp2b)* without heat-shock (E). (G) Quantification of *id1* expression (in the ventricular zone, in the area marked in blue in the upper right-hand corner) displays a significant 1.5-fold increase of *id1* expression upon heat-shock. (H-J’) Immunohistochemistry with antibodies against the RGC marker S100β (blue) and the proliferation marker PCNA (green) of telencephalic transverse sections from WT with heat-shock (H-H’) and *Tg(hs:bmp2b)* without heat-shock (I-I’) and with heat-shock (J-J’) focusing on the ventricular zone of the dorsal telencephalon. White arrows display PCNA+/S100β+ cells. (K) Quantification of the S100β+/PCNA+ cells showing a significant reduction in the heat-shocked (+HS) telencephala compared to the control group. Significance is indicated by asterisks: **, p <0.01; ***, p <0.001. n=3 brains (G), n=15 sections (K). Scale bar = 20 μm (A-C’’, H-J’), 100 μm (D-F). HS: heat-shock.

We next assessed whether BMP2b would affect *id1* expression. Indeed, an increase of *id1* mRNA was triggered by heat-shock in the ventricular zone in comparison to transgenic animals without heat-shock (Figures 2E, F). Heat-shock of a WT adult zebrafish did not elicit induction of *id1* (Figure 2D) demonstrating that the observed response was dependent on the transgene. Quantification of the experimental data revealed that the level of *id1* induction was in the range (Figure 2G) which was previously noted in response to injury (Rodriguez Viales et al., 2015).

We next asked whether the observed induction of *id1* is paralleled by a reduction of proliferating RGCs. For this purpose, sections through the telencephalon of heat-shocked and non-heat-shocked *Tg(hsp70:bmp2b)* animals were labeled with antibodies directed against the RGC marker S100β (März et al., 2010; Zupanc et al., 2005) and the proliferating cell nuclear antigen (PCNA) (Leung et al., 2005; März et al., 2010). In comparison to non-heat-shock controls (Figures 2I, I’), the number of proliferating RGCs was reduced in brains of heat-shocked animals (Figures 2J, J’, white arrows). Upon quantification, the reduction of proliferating RGCs was found to be highly significant (Figure 2K). Heat-shock of WT adult zebrafish did not elicit induction of RGC proliferation (Figures 2H, H’). Taken together, expression of Bmp2b leads to an increase of *id1* expression and a decrease of proliferating RGCs, in line with the notion that BMPs control cell proliferation via activation of *id1*. This pattern of decrease of proliferation and increase of *id1* expression is highly similar to that seen previously as a response to injury of the telencephalon (Rodriguez Viales et al., 2015).

### Inhibition of BMP signaling causes increased proliferation of RGCs

If BMPs were indeed involved in the regulation of RGC proliferation, we would anticipate that blocking BMP signaling decreases *id1* expression and increases proliferation. To test this, we employed the conditional expression of a dominant-negative BMP receptor 1a mutant (Quillien et al., 2011) in which the kinase domain has been deleted. In comparison to non-heat-shocked *Tg(hs:dnBmpr1a)* animals (Figure 3A), a heat-shock resulted in a decrease of *id1* expression in the telencephalon (Figure 3B). Heat-shock of WT animals not carrying the transgene, did not affect *id1* expression (Figure S2A). When quantified on multiple sections, the induced reduction was confirmed to be highly significant (Figure 3C).

**Figure 3.**
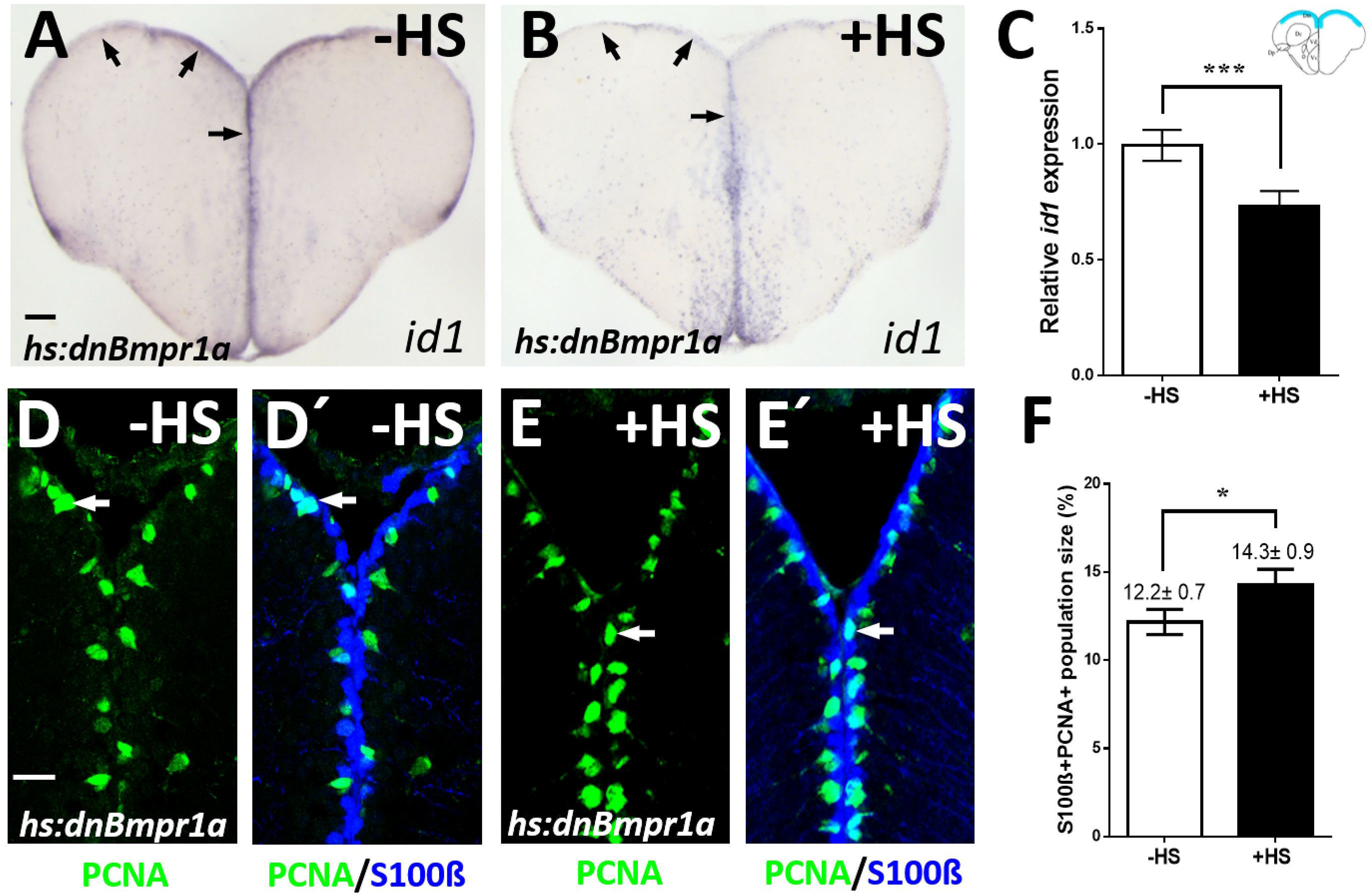
Inhibition of BMP signaling leads to a reduction of *id1* expression and an increase of proliferating RGCs. (A-B) Reduced *id1* expression after inhibition of the BMP pathway by heat-shock of *Tg(hs:dnBmpr1a)* animals. Black arrows point at the expression of *id1* in the ventricular zone of *Tg(hs:dnBmpr1a)* animals with heat-shock (A) and without heat-shock (B). (C) Quantification of *id1* expression of brains shown in A-B (quantified region is marked in blue in the scheme in the upper right-hand corner) indicates that inhibition of BMP signaling leads to a significant reduction of *id1* expression. (D-E’) Immunohistochemistry of telencephalic transverse sections for RGCs (S100β, blue) and proliferating cells (PCNA, green), focusing on the ventricular zone of the dorsal telencephalon. White arrows indicate PCNA+/S100β+ cells. (F) Quantification of the population size of S100β+/PCNA+ cells exhibiting a significant increase in heat-shocked compared to non-heat-shocked telencephala. Significance is indicated by asterisks: *, 0.01≤ p <0.05; ***, p <0.001. n=3 brains (C), n=15 sections (F). Scale bar = 20 μm (D-E’), 100 μm (A-B). HS: heat-shock.

We next asked whether this induced reduction of *id1* expression is accompanied by an increase of proliferation of RGCs. In comparison to non-heat-shocked controls (Figures 3D, D’), the *hs:dnBmpr1a* expressing animals showed an increase of PCNA+/S100β+ (white arrows) and thus proliferating RGCs (Figures 3E, E’ and F). Taken together, these data demonstrate that inhibition of BMP signaling by employment of the dominant-negative receptor, leads to the reverse effect of what was observed upon forced expression of BMP2b. This also suggests that the lack of pSmad staining in RGCs of wild-type animals (Figure 2A’) is due to limited sensitivity of the assay. The proliferation of neural stem cells in the zebrafish telencephalon thus appears to be controlled by BMPs. The observed effects on *id1* expression are consistent with our preliminary results indicating a role of *id1* as a repressor of RGC proliferation in response to injury (Rodriguez Viales et al., 2015). In conclusion, the presented data show that BMPs regulate *id1* expression and most likely influence cell proliferation by acting through *id1*.

### BMP and Notch signaling cooperate in controlling RGC quiescence

Notch3 and the Notch-target gene *her4.1* have previously been proposed to be mainly expressed in quiescent RGCs (Alunni et al., 2013; Than-Trong et al., 2020). In agreement with its expression pattern, Notch3 negatively controls the proliferation of the RGCs (Alunni et al., 2013). It was also previously suggested that there are distinct classes of RGC populations with different proliferative potential (Cosacak et al., 2019; Diotel et al., 2010; Diotel et al., 2016; Lange et al., 2020; März et al., 2010; Than-Trong et al., 2020). Our observation that BMPs mediate RGC quiescence raises the question of how these two pathways cooperate.

Since they may operate in distinct RGC populations, we first assessed whether *id1* and *notch3* or *her4.1* are expressed in the same RGCs (Figures 4A, B, Figure S3A-C’’). We hybridized sections through the telencephalon of *Tg(id1-CRM2:gfp)* fish with a *notch3* antisense RNA probe and counted expressing and co-expressing cells (Figure 4A, Figures S3A-A’’). In total, co-expression was detected in 83.7% cells suggesting that Notch3 and BMP signaling target the same RGC populations in vast majority (Figure 4A). To assess co-expression of *her4.1* and *id1*, we crossed *Tg(her4.1:gfp)* transgenic fish (Yeo et al., 2007) into a *Tg(id1-CRM2:mCherry)* background (Zhang et al., 2020). We counted 88.7% cells co-expressing the two transgenes (Figure 4B, Figures S3B-C’’’) confirming the result of the *notch3/id1* co-expression analysis. This suggests that the two regulators of stem cell quiescence are largely co-expressed and therefore act in most cases in the same cells. Curiously, we found significant, but much smaller fractions of RGCs that expressed only *Tg(her4.1:gfp)* or *Tg(id1-CRM2:mCherry).* We assessed whether there are differences in proliferative potential between the *her4.1+/id1+* RGCs and the RGCs expressing only one of the two genes. Indeed, 31.7% RGCs expressing *Tg(her4.1:gfp)* and 56.7% RGCs expressing *Tg(id1-CRM2.:mCherry)* only were PCNA+, while only 4.7% RGCs positive for both markers were also PCNA+ (Figure S3D). This suggests that cells expressing either *id1* or *her4.1* are more frequently cycling than the double positive cells.

**Figure 4.**
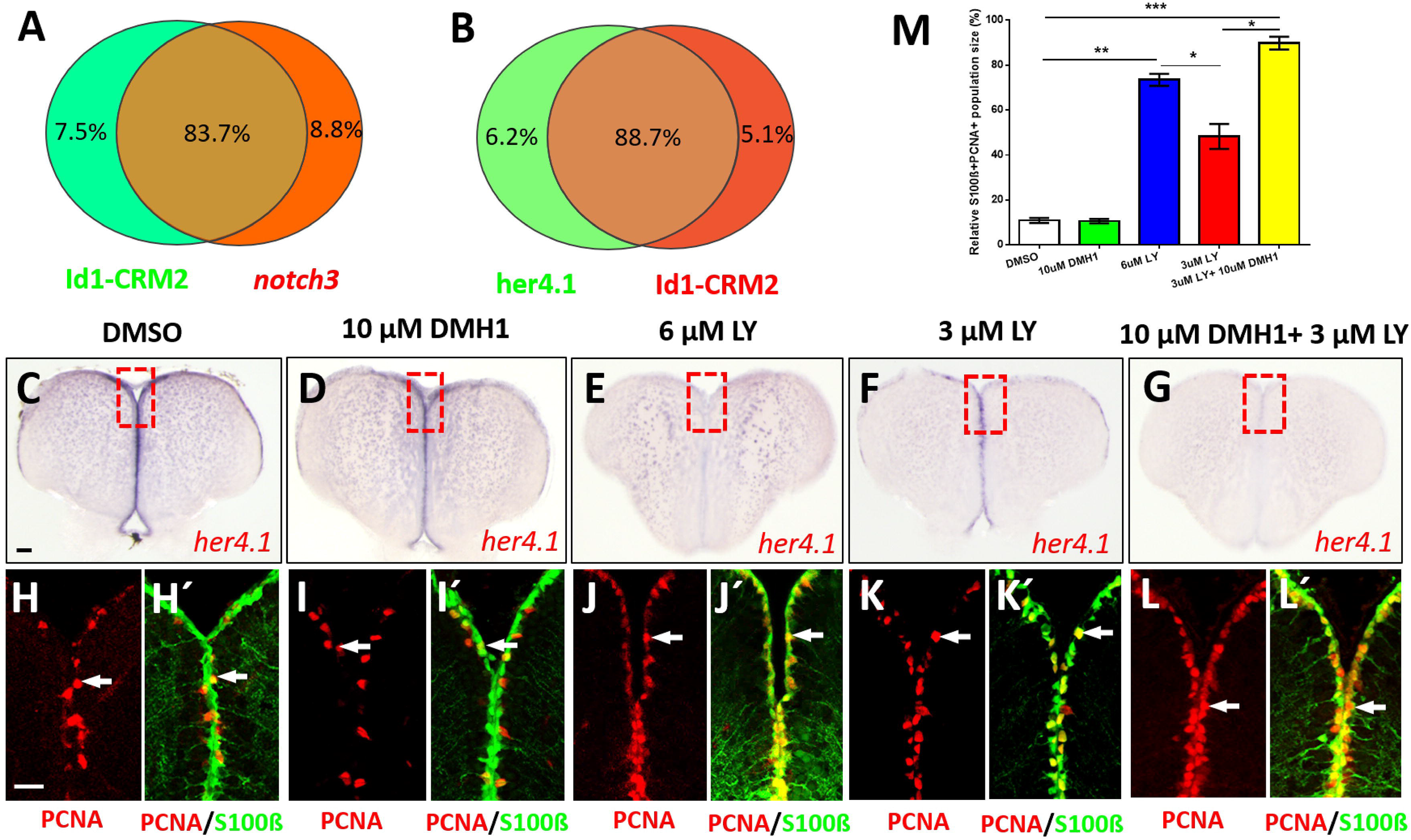
Notch and BMP signaling pathways interact to control quiescence of NSCs. (A-B) Summary of co-expression analyses of *Tg(id1-CRM2-mCherry)* with *notch3* (A) and with *Tg(her4.1:gfp)* (B). RGCs express *id1* and the Notch signaling components Notch3 and *her4.1.* (For examples of sections see Figure S3A-C’’’). (C-G) Expression of *her4.1* revealed by ISH on control telencephala (DMSO) or telencephala treated with different concentrations of LY411575 (LY), or a combination of LY and DMH1. (D) 10 μM DMH1 does not influence the expression of *her4.1* strongly. (E) Notch blockage reduces *her4.1* expression along the ventricular zone. (F) After reduction of the concentration of LY to 3 μM, *her4.1* expression is still recognizable in the ventricular zone. (G) Combination of 3 μM LY with 10 μM DMH1 blocks *her4.1* expression in the ventricular zone suggesting that the two pathways interact. Red rectangles (C-G) illustrate regions of immunostaining in H-L’. (H-L’) Cross-sections of the pallial ventricular zone following different concentrations of LY treatment, immunostained for the RGC marker S100β (green) and the proliferation marker PCNA (red). The proportion of S100β+/PCNA+ cells is increased in the groups treated with 6 μM LY alone (J-J’) or treated with a combination of 3 μM LY and 10 μM DMH1 (L-L’). White arrows indicate PCNA+/S100β+ cells. (M) Quantification of the relative population size of S100β+/PCNA+ cells under different conditions. Significance is indicated by asterisks: *, 0.01≤ *p* <0.05; **, *p* <0.01; ***, *p* <0.001. *n*=3 brains. Scale bars: 20 μm (H-L’), 100 μm (C-G).

Previously, we showed that *id1* expression in the adult telencephalon is not affected by the Notch inhibitor LY ((Rodriguez Viales et al., 2015) and Figure S4)) demonstrating that *id1* expression and hence also BMP signaling is independent of Notch signaling. Reversely, we asked now what happens to expression of *her4.1*, if either one or both pathways are inhibited by chemical interference. Treatment with 10 μM DMH1, an inhibitor of the canonical BMP signaling pathway (Hao et al., 2010), affected the expression of *her4.1* marginally (Figure 4D). Application of 6 μM LY411575 (LY), a gamma-secretase inhibitor and potent Notch signaling blocker (Fauq et al., 2007), eliminated expression of endogenous *her4.1* entirely (Figure 4E), while 3 μM LY (Figure 4F) caused a reduction of *her4.1* expression relative to DMSO treated controls (Figure 4C). Co-application of 10 μM BMP inhibitor DMH1 together with 3 μM Notch inhibitor LY completely abolished *her4.1* expression (Figure 4G). Thus, BMP signaling has a positive effect on *her4.1* expression suggesting that the two pathways converge on this transcription factor.

Next, we asked whether also cell proliferation is affected by chemically blocking the Notch and BMP pathways. We co-stained telencephalic sections of inhibitor-treated animals with antibodies against PCNA and S100β (Figures 4H-L’). While exposure to 10 μM DMH1 alone had no effect on the number of PCNA+ RGCs (white arrows) (Figures 4H-I’, M) addition of 6 μM LY (Figures 4H-H’, J-J’, M) or 3 μM LY (Figures 4H-H’, K-K’, M) caused an increase in proliferating cells. When suboptimal inhibitor concentrations (3 μM LY, 10 μM DHM1) were co-applied (Figure 4G), a more than additive increase in the number of proliferating cells (white arrows) was scored (Figures 4H-H’, L-L’, M). This increase of proliferating RGCs in co-inhibited animals is similar to the effect on *her4.1* expression. These observations further support the notion that BMP and Notch signaling pathways converge on *her4.1*.

Inhibitor experiments are limited by the efficacy of the inhibitors. To confirm the suggested effect of BMP signaling on *her4.1* expression, we analyzed *her4.1* expression in transgenic *Tg(hs:dnBmpr1a)* animals expressing the dominant negative BMP receptor in response to heat-shock. When exposed to elevated temperature, *her4.1* expression was reduced in comparison to transgenic animals not exposed to heat shock (Figure S5). Thus, *her4.1* appears to be under the control of BMP signaling consistent with our model that *her4.1* expression being a convergence point of the BMP and Notch pathways.

### Genetic removal of *id1* activity leads to increased proliferation in the telencephalon

Previously, transient reduction of Id1 activity by cerebroventricular microinjection of a vivo-morpholino led to increased proliferation of RGCs (Rodriguez Viales et al., 2015). Given the temporal and spatial limitation of this method and its potential toxic impact on the transfected tissue (Ferguson et al., 2014), we wished to confirm the results from these transient studies with a stable genetic approach. Moreover, our pharmacological inhibitor experiments suggested that BMPs control *her4.1* expression. If this effect on *her4.1* is mediated through *id1*, loss-of-function of *id1* should result in a reduction of *her4.1* expression.

To test these hypotheses, we generated a knock-out of *id1* using a CRISPR-Cas9 approach. The identified mutant contains a 2 bp deletion in the start codon of the *id1* coding sequence, which leads to a non-functional shorter Id1 protein (Figure S6A). Homozygous *id1*^*ka706/ka706*^ animals were viable and reached adulthood without gross morphological defects. This suggests that other highly related *id* genes present in the zebrafish genome (Diotel et al., 2015a) may compensate for the loss of a possible function of *id1* during development.

To assess the proposed role of *id1* in the regulation of proliferation of RGCs, we stained telencephalic cross-sections of 6 month old WT (Figures 5A-A’’) and *id1*^*ka706/ka706*^ animals (Figures 5B-B’’) with the RGC marker S100β and the proliferation marker PCNA. A two-fold increase in the number of PCNA+ cells (yellow arrows) could be detected in the ventricular zone of the mutants in comparison to that of WT siblings (Figure 5C). When the number of PCNA+ / S100β+ cells (white arrows) was compared between mutant and WT, a similar increase was noted (Figure 5D), suggesting that in *the id1*^*ka706/ka706*^ mutant about twice as many RGCs are in a proliferative state. We also tested whether the increased number of proliferating RGCs has an effect on *ascl1a* expression which has been implicated in proliferation as well as commitment to neurogenesis in mammals (Andersen et al., 2014; Blomfield et al., 2019; Castro et al., 2011). To this end, we stained transverse sections through WT and *id1*^*ka706/ka706*^ telencephala with an antisense probe directed against the neural precursor gene *ascl1a* (Figures 5E-F). A 1.5-fold increase of *ascl1a* expression was observed in the mutant telencephalon (Figure 5G) which was verified by quantitative PCR (qPCR; (Figure 5H)).

**Figure 5.**
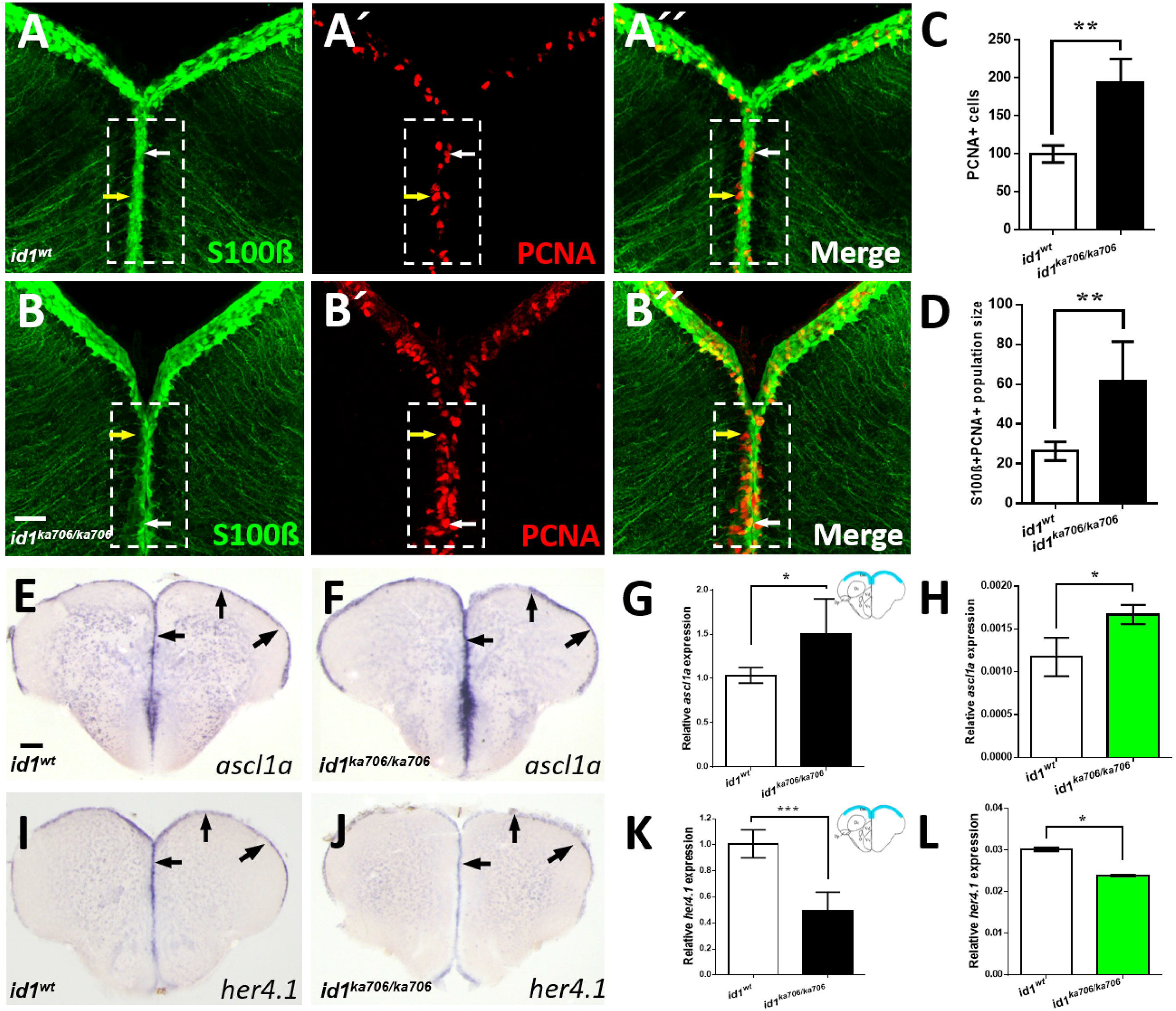
Loss of *id1* function leads to an increased number of proliferating RGCs. (A-B’’) RGCs marked by immunohistochemistry with S100β (green) and PCNA (red) antibodies on telencephalic cross-sections from adult *id1*^*ka706/ka706*^ and WT siblings. The number of PCNA+ cells per brain is increased in the mutant (B’) compared to WT control brains (A’). White arrows show PCNA+/S100β+ cells, yellow arrows show PCNA+ cells. (C) Relative population size of PCNA+ cells in WT and *id1*^*ka706/ka706*^ brains. (D) Quantification of the number of proliferating RGCs (type II cells, S100β+/PCNA+ cells) in *id1*^*ka706/ka706*^ and WT siblings. (E-F) Expression of *achaete-scute-like1a (ascl1a)* mRNA is increased in *id1*^*ka706/ka706*^ telencephala. Black arrows show the expression of *ascl1a* in the ventricular zone of *id1*^*ka706/ka706*^ mutants and WT siblings. (G) Quantification of *ascl1a* expression (scheme in the upper right-hand corner displays the quantified area in blue). (H) RT-qPCR quantification confirms induction of *ascl1a* in *id1*^*ka706/ka706*^. (I-J) Expression of *her4.1* mRNA is reduced in *id1*^*ka706/ka706*^ telencephala. Black arrows show the region of expression of *her4.1* in the ventricular zone of *id1*^*ka706/ka706*^ mutants and WT siblings. (K) Quantification of *her4.1* expression (scheme in the upper right-hand corner displays the quantified area in blue). (L) RT-qPCR quantification confirms reduction of *her4.1* in *id1*^*ka706/ka706*^ telencephala. Significance is indicated by asterisks: *, 0.01≤ *p* <0.05; **, *p* <0.01. *n*=3 brains (A-D), *n*=15 sections (G, K), n=5 telencephala (H, L). Scale bars: 20 μm (A-B’’) 100 μm (E-F, I-J).

To confirm the suggested effect of BMP/Id1 signaling on *her4.1* expression, we analyzed *her4.1* expression in *id1*^*ka706/ka706*^ mutant animals. Transverse sections through WT and *id1*^*ka706/ka706*^ telencephala stained with an antisense probe directed against the *her4.1* gene (Figures 5I-J) show a reduction of *her4.1* expression in the *id1*^*ka706/ka706*^ mutant compared to WT animals (Figures 5I-K). This reduction was validated by qPCR (Figure 5L).

These data confirm our preliminary results from vivo-morpholino knock-down experiments (Rodriguez Viales et al., 2015): *id1* plays a role in the control of stem cell quiescence. Moreover, *id1* is required for expression of *her4.1* in RGCs supporting the notion that *her4.1* is an integration point of BMP and Notch signaling in the control of RGC proliferation.

Next, we asked whether the higher number of activated NSCs in the mutant would lead to an increase in the number of RGCs in the mutant. We found a slightly but significantly increased number of RGCs in the mutant telencephala (Figure S6B). Thus, the pool of stem cells is not depleted despite the increased neurogenesis in the *id1* mutant. This suggests that mechanisms remain in operation that maintain the stem cell pool in the mutant under normal physiological conditions.

### *id1* is essential to preserve long term maintenance of the regenerative capacity in the injured telencephalon

The level of *id1* expression in individual cells, as well as the total number of cells is increased five days after inflicting a lesion to the telencephalon. This response of *id1* expression followed the induction of RGC proliferation (Rodriguez Viales et al., 2015). We speculated that Id1 is required to dampen the proliferative response to prevent exhaustion of the stem cell pool in the regenerating brain. We addressed this hypothesis here by causing stab wounds in the telencephalon of *id1*^*ka706/ka706*^ and WT control animals. In the wounding protocol, the right hemisphere of the telencephalon is injured by introducing a needle through the skull while the left hemisphere remains uninjured (März et al., 2011; Schmidt et al., 2014). Afterwards, the animals were allowed to recover 1 month before a second lesion was inflicted (Figure 6A). Stab wounded animals were sacrificed 5 days after the first or the second lesion or kept for another 3 months (Figure 6A). For examination of proliferation and repair of lesion, sections through the telencephalon were co-stained with antibodies against the proliferation marker PCNA and the RGC marker S100β. As expected from the previous experiments, mutant animals showed a stronger proliferative response of the RGCs than the WT control, 5 days after the first lesion (Compare Figures 6C, C’ with Figures 6B, B’, white arrows). This is in striking contrast to the situation after the second injury: While WT siblings had many PCNA+ RGCs and thus a profound proliferative response to the second lesion, the mutants displayed less proliferating RGCs (Figures 6E-G). Moreover, when examined 3 months after the second wounding, the WT animals (n=16) had all healed the second wound (Figures 6H, I) and only 5 animals showed some mild traces of the injury (Figure 6I), In contrast, 7 of the 11 mutant animals showed massive tissue defects (Figure 6J) and only 4 had repaired the second lesion partially with clear signs of the lesion remaining (Figure 6K, white arrows). Also, the survival rate was slightly less in the mutants at 3 months (Table S2).

**Figure 6.**
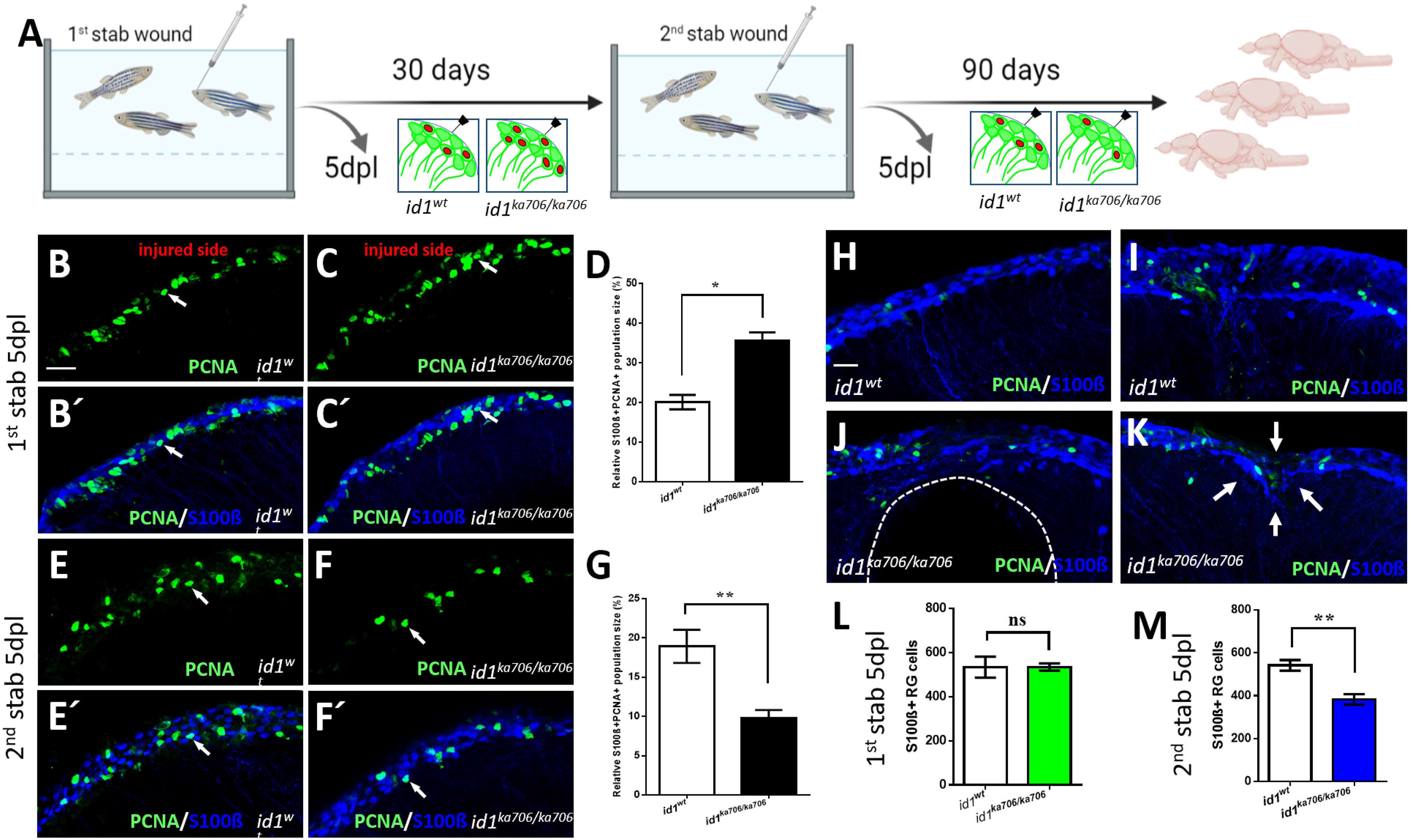
*id1* is required to maintain the stem cell pool and regenerative capacity of the telencephalon. (A) Experimental layout: 6 month old adult *id1*^*ka706/ka706*^ fish (n=25) and WT siblings (n=25) were stabbed with a needle into the right hemisphere of the telencephalon. Fish (n=2) of each group were sacrificed at 5 days post lesion (dpl) and the remaining fish were allowed to recover for one month before a second stab wound was inflicted. Five days after the second lesion, fish (n=7) from each group were sacrificed for analysis. The remaining fish were kept for another 3 months before they were sacrificed (n=16 for WT and n=11 for *id1*^*ka706/ka706*^ mutants). Note that five mutant fish showed signs of suffering after the second wounding and were sacrificed before the end of the three months recovery. (B-C’, E-F’, H-K) Double immunohistochemistry with S100β (blue) and PCNA (green) antibodies was carried out to mark proliferating RGCs on telencephalic cross-sections from *id1*^*ka706/ka706*^ mutants and WT siblings at the different time points. (B-C’) After the first stab, *id1*^*ka706/ka706*^ mutants showed a higher number of S100β+/PCNA+ cells (white arrows). (D) Quantification of the population size of PCNA+/S100β+ cells in relation to the total number of RGCs (S100β+ cells) in WT and *id1*^*ka706/ka706*^ telencephalic cross-sections after the first stab wound. (E-F’) After the second injury, *id1*^*ka706/706*^ mutants showed less PCNA+/S100β+ cells (white arrows) compared to WT siblings. (G) Quantification of the population size of PCNA+/S100β+ cells in relation to the total number of RGCs in cross-sections through WT and *id1*^*ka706/ka706*^ telencephala. (H-K) WT fish had repaired the lesion in the telencephalon 3 months after the second stab without signs of the injury (H) or with only mild signs of slight tissue disorganization (I). In contrast, *id1*^*ka706/ka706*^ mutants showed severe tissue lesions (J), such as a hole in the parenchyma of the telencephalon (dashed line) or dents in the parenchyma with tissue disorganization (K, white arrows). (L-M) Quantification of the RGCs (S100β+ cells) in the injured side after inflicting the first (L) and second (M) stab wound in WT and *id1*^*ka706/ka706*^ mutants showing a stepwise reduction of the number of RGCs. (Compare also with the increased number of RGCs in the uninjured mutant (Figure S6B)). Significance is indicated by asterisks: ns, not significant; *, 0.01≤ *p* <0.05; **, *p* <0.01. Scale bars: 20 μm (B-C’, E-F’, H-K).

We next asked whether the overall number of RGCs in the telencephalon decreased in the mutant. The number of RGCs (S100β+ cells) was comparable 5 days after the first lesion between the mutant and WT telencephala (Figure 6L). Note that uninjured animals showed a slightly increased number of RGCs (Figure S6B). After the second injury, the number of RGCs had dropped by 32% in the mutants (Figure 6M). Hence, lack of *id1* function causes not only loss of proliferation of RGCs but also shrinkage of the RGC pool as a whole in the injured animal. This suggests that an injury leads to an imbalance towards neurogenesis that cannot be controlled by the mechanisms of stem cell maintenance in operation in parallel to the BMP/Id1 regulatory circuit.

In summary, in absence of *id1*, the ventricular zone initially displayed a much more profound proliferative response to a lesion (Figure 7). Upon second injury, however, proliferating RGCs were severely reduced in the mutant. This, together with the overall loss of RGCs, suggests depletion of proliferation-competent stem cells. These data show that *id1* is required to maintain the stem cell pool after wounding and thus the long-term capacity to efficiently repair wounds in the telencephalon.

**Figure 7:**
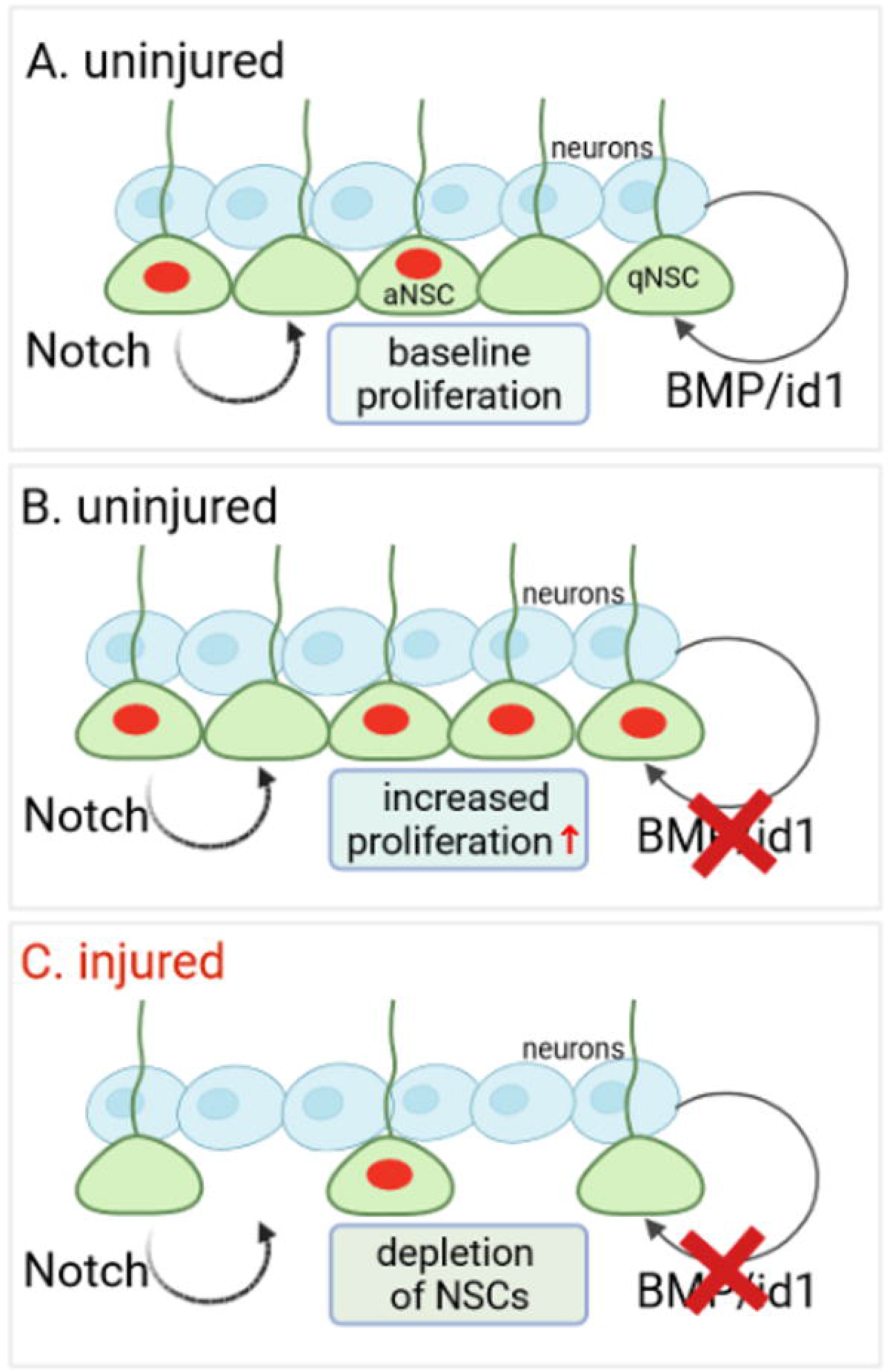
Model of regulation of NSC proliferation by Notch and BMP/Id1 signaling. Constitutive neurogenesis in WT telencephala controlled by cooperation of Notch signaling originating from NSCs in the telencephalic ventricle and BMP signaling stemming from neurons close to the NSCs. Activated NSCs/RGCs (aNSC) are indicated by red nuclei. (A) Constitutive neurogenesis in *id1*^*ka706/ka706*^ mutant telencephala where BMP/Id1 signaling is not active. There is an increased number of proliferating NSCs (red nuclei).. (C) Regenerative neurogenesis in *id1*^*ka706/ka706*^ mutant telencephala (after injury). The NSC pool becomes depleted because of ongoing proliferation of NSCs.

## Discussion

Zebrafish have a remarkable ability to generate new neurons and to heal injuries in the central nervous system. Here, we show that, in the telencephalon, the basis of this long-term neurogenic potential involves the management of the stem cell pool by the BMP/Id1 signaling pathway. Interference with BMP signaling led to modulation of *id1* expression which in turn repressed proliferation of NSCs/RGCs. Genetic ablation of *id1* activity initially caused increased proliferation of NSCs and then depletion of the stem cell pool and failure to repair repeated wounding of the telencephalon. Our data are consistent with a model in which the regenerating nervous tissue feeds back on the stem cell pool via BMP/Id1 signaling thereby promoting quiescence after the initial rise of proliferation in response to injury. Cross-talk of BMP/Id1 signaling with the Notch signaling pathway appears to occur at the level of expression of the transcription factor *her4.1*. In the absence of *id1* and under homeostatic conditions of constitutive neurogenesis, Notch signaling appeared to be sufficient to maintain the stem cell pool. However, when challenged by repair of injury, this safeguard is not sufficient anymore and the NSC pool is depleted in the *id1* mutant. Thus, the NSC pool is maintained by two-tiered regulatory mechanisms involving both Notch and BMP/Id1 signaling.

### Neuron-derived BMPs control proliferation of NSCs

Several lines of evidence demonstrate that BMPs control proliferation of RGCs. Conditional expression of BMP2b caused reduction of RGC proliferation while blocking expression of BMP signaling via expression of dnBMPR1a led to increased proliferation of RGCs. Pharmacological inhibition of BMP signaling had the same effect as expression of the dominant negative BMP receptor, reinforcing our conclusions.

The source of BMPs in vicinity to the RGCs is the neurons in the telencephalic parenchyma. In contrast to stem cells in the mouse telencephalon (Lim et al., 2000), RGCs of the zebrafish telencephalon do not express BMPs. Single cell-sequencing analysis suggests, however, that RGCs are fully competent to respond to BMP signals expressing all the key members of the intracellular BMP signaling cascade. Moreover, when BMPs are over-expressed, only RGCs show detectable pSmad staining in cross-sections through the entire parenchyma of the telencephalon. These data suggest that neurons are the BMP emitting cells and RGCs are the BMP receiving cells.

*id1*, whose expression is increased by wounding in a delayed fashion and following the activation of proliferation of RGCs (Rodriguez Viales et al., 2015), depends on BMP signaling. *id1* expression in RGCs including its up-regulation in response to injury is faithfully mimicked by a *cis-*regulatory module derived from the *id1* gene. This module requires Smad1/5/9 DNA binding elements to function (Zhang et al., 2020) in support of the notion that *id1* is a direct downstream target of BMP signaling.

A noteworthy phenomenon is the restricted activation of proliferation in the injured hemisphere only (März et al., 2011; Rodriguez Viales et al., 2015). Also, activation of *id1* expression occurs only in the injured hemisphere (Rodriguez Viales et al., 2015; Zhang et al., 2020). This suggests that not only induction of proliferation by injury but also its dampening by BMPs back to the baseline levels of constitutive neurogenesis is restricted to the injured hemisphere. It is thus unlikely that BMPs are released into the cerebrospinal fluid as they would otherwise also affect RGCs in the uninjured hemisphere. The long processes of RGCs spanning the entire parenchyma may play a role as sensors of BMP signals. It remains to be determined whether BMP expression is increased in response to repair of the tissue relative to normal levels of expression. By both RNA ISH and antibody staining, we did not detect changes of expression of the four tested *bmp* genes or the BMP2b protein (unpublished). This may be a matter of sensitivity or BMPs are regulated at the level of processing or other post-translational mechanisms that escaped our detection due to lack of tools. Irrespectively, our data strongly suggest that neurons are the BMP producing and RGCs are the BMP sensing cells. This cross-talk is somehow up-regulated by regeneration leading to dampening of proliferation. Our data are consistent with a model in which the nervous system maintains, via BMP signaling, a feed-back loop between the regenerating nervous tissue and the stem cell niche thereby adapting the proliferative rate of the stem cells to the needs of tissue homeostasis and regeneration.

### Id1 mediates long-term maintenance of the regenerative capacity

Mutation of *id1* leads to an increased number of proliferating RGCs and elevated formation of neuroblasts committed to a neural fate. These changes are similar in magnitude to those observed previously in response to injury of the telencephalon (Rodriguez Viales et al., 2015; Zhang et al., 2020). RGCs divide by symmetric and asymmetric divisions (Barbosa et al., 2015; Rothenaigner et al., 2011). Mechanisms have to be in place that prevent depletion of the stem cell pool, especially when production of new neurons was boosted by injury. Our analysis of the *id1* mutant as well as the BMP gain- and loss-of-function experiments suggest exactly such a role for the BMP/Id1 signaling system. BMP/Id1 signaling promotes a non-proliferative state and thereby maintenance of the NSC pool.

After the first injury of the telencephalon, the *id1* mutants showed a stronger proliferative response than the WT siblings. When wounding was repeated, the mutants presented a significantly reduced proliferative capacity after the second round of injury. This suggests that the stem cell niche in *id1* mutants was significantly deprived of proliferation-competent RGCs after the first injury/regeneration cycle. This is reflected by a reduction of overall RGC counts after wounding. Fully in agreement with this loss of RGCs, *id1* mutants failed to repair the second lesion in contrast to WT siblings treated in the same way. *id1,* and consequently also BMP signaling, are thus required by the adult telencephalon to maintain its capacity of regenerative neurogenesis. This shift back to baseline proliferation could be mediated by two distinct mechanisms. BMP/Id1 signaling could prevent quiescent type I RGCs from entering the cell cycle or it could actively promote return of proliferation competent type II RGCs to return to quiescence after injury-induced rise of proliferation.

### A conserved function of BMP signaling in stem cell maintenance

It is striking that the CRM2 driving *id1* expression (Zhang et al., 2020) in RGCs of the telencephalon is conserved between mammals and zebrafish. Even more astonishing is that the human homologue of CRM2 mediates expression in the telencephalon of the zebrafish in a pattern identical to that of the endogenous element (Zhang et al., 2020). Moreover, the function of the *id1-*CRM2 depends on the integrity of conserved BMP response elements. This suggests that the basic mechanisms of BMP signaling are conserved and presumably also in operation in the two distantly related vertebrate species.

In rodents, multiple functions of BMP signaling ranging from control of proliferation, differentiation and gliogenesis to quiescence of NSCs have been reported in various contexts (see (Choe et al., 2015) for review). Fetal NSCs respond to BMPs with proliferation (Panchision and McKay, 2002). In contrast, blockage of BMP signaling in the dentate gyrus suggests that BMPs mediate quiescence (Mira et al., 2010) in agreement with our findings in the zebrafish. In the mouse SVZ, this control of quiescence involves locally balanced BMP levels via expression of the BMP clearance receptor LRP2 in ependymal cells. This control appears to be restricted to the SVZ as the subgranular zone of the mouse telencephalon does not express LRP2 (Gajera et al., 2010). In contrast to zebrafish NSCs, in the murine SVZ, NSCs (type B cells) express BMP2 and 4 (Lim et al., 2000) pointing at significant differences between zebrafish. Moreover, conditionally blocking Smad4 in Type B cells of the SVZ leads to oligodendrogenesis (Colak et al., 2008). We did not observe this shift in the zebrafish telencephalon but rather noted an increase in neurogenesis when we blocked the BMP/Id1 signaling axis. Murine type B stem cells of the SVZ show activation of Smads via phosphorylation and also ID1 is expressed at high levels in these cells (Nam and Benezra, 2009). Indirect evidence also points to involvement of BMPs in regulation of proliferation in the SGZ in the hippocampus of mice (Choe et al., 2015). The BMP/Id1 regulatory module is evolutionary ancient and, like Notch signaling, it has been deployed in multiple processes during development and body homeostasis (Wang et al., 2014). The data from studies in the mouse telencephalon suggest that the BMP/Id1 regulatory module is also employed in the control of quiescence in NSCs of the murine telencephalon, even though mice do not show a strong regenerative capacity.

### Cooperation between BMP and Notch pathways in the control of stem cell quiescence

Multiple lines of previous evidence suggest that Notch signaling controls proliferation of RGCs in the zebrafish telencephalon (Alunni et al., 2013; Chapouton et al., 2010; de Oliveira-Carlos et al., 2013; Than-Trong et al., 2018). In particular, Notch3 has been implicated in the control of RGC quiescence (Alunni et al., 2013). Notch signaling appears to act on short-range between NSCs and activated neural precursors (type III cells) (Dray et al., 2021a). This process bears resemblance to lateral inhibition initially proposed to select the precursor cell of sensory bristles in the Drosophila cuticle (Simpson, 1990). Thus, the dynamics of the NSCs generate an intrinsic cue that assures their long-term maintenance (Dray et al., 2021a). Our data here suggest that intrinsic Notch signaling is not the only mechanism that safeguards maintenance of NSCs in the zebrafish telencephalon but also BMP mediated cross-talk between neurons and the stem cells.

When Notch signaling is blocked pharmaceutically, *her4.1* expression is reduced and proliferation is increased (Alunni et al., 2013). Strikingly *her4.1* expression is also affected when BMP signaling or *id1* activity is altered: Lack of *id1* function leads to a reduction of *her4.1* expression. Moreover, pharmacological treatment with sub-optimal concentrations of BMP and Notch inhibitors results in cooperative effects with loss of *her4.1* expression. However, inhibition of Notch signaling does not seem to influence *id1* expression ((Rodriguez Viales et al., 2015), this manuscript). Our data suggest that BMP signaling acts in parallel of and converges with Notch signaling on the expression of *her4.1*. A similar interaction of BMP and Notch signaling was recently uncovered in the control of angiogenesis in zebrafish embryos (Esser et al., 2018). It remains to be shown, however whether the control of quiescence is mediated entirely through Her4.1 or whether other mechanisms acting in parallel exist.

It is unclear how Id1 and Her4.1 affect cell proliferation. During mouse neurogenesis and myogenesis, the Notch effector Hes1, related to zebrafish Her4.1, is either expressed at high levels or oscillates (Sueda et al., 2019; Sueda and Kageyama, 2020). The downstream proneural gene *ascl1* is completely suppressed in quiescent stem cells where the repressor Hes1 is expressed at high levels. Expression of the regulator *Hes1* can start oscillating due to an autoinhibitory loop and, as a consequence, the expression of *ascl1* becomes also oscillatory. The oscillatory state allows escape from cell-cycle arrest (Sueda et al., 2019; Sueda and Kageyama, 2020). The HLH protein Id1 is a negative regulator of bHLH proteins like Her4.1 and Hes1 (Roschger and Cabrele, 2017). Moreover, Id1 binds to her4.1 *in vitro* (Rodriguez Viales et al., 2015). Disrupting the postulated negative feedback loop of her4.1/Hes1 would increase expression of *her4.1/hes1*. In agreement, blocking BMP signaling or knocking-out *id1* resulted in reduction of *her4.1* expression in the zebrafish telencephalon. It remains to be seen whether *ascl1* is indeed a key regulator of proliferation in the zebrafish telencephalon.

### Why are two pathways necessary to control stem cell quiescence?

A question is why are two pathways necessary to control proliferation of NSCs. Previously, it has been suggested that distinct RGCs exist with respect to marker expression and cell cycle progression along the ventricular zone of the RGCs (Cosacak et al., 2019; März et al., 2010). A simple explanation may thus be that the distinct populations employ different mechanisms to control stem cell quiescence. However, the observation that most stem cells co-express *id1* and *her4.1* suggests that the majority of RGCs receives input from both signaling systems. The large number of RGCs expressing both *her4.1* and *id1* may be the large fraction of deep quiescent cells postulated previously (Than-Trong et al., 2020). In agreement, when we checked for expression of the proliferation marker PCNA, the *id1/her4.1* double-positive cells expressed PCNA less frequently than the single *id1* or *her4.1* expressing cells. It is, however, also possible that these *id1+/her4.1-/*PCNA+ and the *id1-/her4.1+/*PCNA+ cells are transition state cells that are on the way back to quiescence or in transition to neuroblasts (Type IIIa cells (März et al., 2010) also called activated neural progenitors (aNPs) (Dray et al., 2021a)).

Clearly, Notch appears to be almost sufficient to maintain a normal stem cell pool in the uninjured *id1* mutant. RGC proliferation is increased in the *id1* mutant. However, Notch maintains under these conditions a high number of RGCs, - even slightly higher than what was found in WT siblings. The stem cell population shrank only after injury in the *id1* mutant.

This clearly allocates distinct functions to Notch and BMP/Id1 signaling. While Notch3 signaling appears to have a predominantly homeostatic function during constitutive neurogenesis, BMP/Id1 comes into play when the proliferation control system is moved out of balance by injury. Under these circumstances, BMP/Id1 is necessary to prevent depletion of the stem cell pool and eventual loss of the regenerative capacity. Taken together, maintenance of stem cells by BMP/Id1 signaling is a key mechanism that underlies the remarkable ability of adult zebrafish to heal even severe injuries of the forebrain. Clearly our data suggest that stem cells are not only maintained by niche-intrinsic cues but also via neuron/radial glial communication.

## ACKNOWLEDGMENTS

We thank Nadine Borel and the fish facility staff for fish care. Anne Schröck and Elisa Kämmer for technical support. Ferenc Muller (University of Birmingham), Laure Bally-Cuif (Institut Pasteur) and David Morizet (Institut Pasteur) for comments, suggestions and proofreading of the manuscript. Elise Cau (CBD, Toulouse), Matthias Hammerschmidt (University of Cologne) and Jeroen Bakkers (Hubrecht Institute) for sharing zebrafish transgenic lines and constructs. We are grateful for support by the EU IP ZF-Health (Grant number: FP7-242048), the Deutsche Forschungsgemeinschaft (GRK2039), the program BioInterfaces in Technology and Medicine of the Helmholtz Association and the European Union’s Horizon 3952020 research and innovation program under the Marie Sklodowska-Curie grant agreement No. 643062 (ZENCODE-ITN).

## AUTHOR CONTRIBUTIONS

S.R., and U.S: designed the experiments and supervised the work, analyzed the data, and wrote the manuscript; G.Z, L.L and T.B: conducted the experiments and analyzed the data; N.D and M.T: analyzed the data; F.C : performed the single RNA sequencing data analysis; G. Z and M.T., analyzed and quantified the data. All authors contributed to the article and approved the submitted version.

## Materials and Methods

### Zebrafish strains and husbandry

Experiments were performed on 6-12 month old AB wild-type (WT), *Tg(id1-CRM2:GFP)* (Zhang et al., 2020), *Tg(her4.1:GFP); Tg(id1-CRM2:mCherry)* (Yeo et al., 2007) and Zhang et al., 2020 respectively), *Tg(olig2:gfp)* (Park et al., 2007), *Tg(hs:bmp2b)* (Chocron et al., 2007), *Tg(hs:dnBmpr1a)* (Quillien et al., 2011) and *id1*^*ka706/ka706*^ (this study) zebrafish. Zebrafish housing and husbandry were performed following the recommendations by (Alestrom et al., 2019). All animal experiments were carried out in accordance with the German protection standards and were approved by the Government of Baden-Württemberg, Regierungspräsidium Karlsruhe, Germany (Aktenzeichen 35-9185.81/G-288/18).

### Stab wound, chemical treatment and heat shock of adult zebrafish

Stab wounding was performed as described (März et al., 2011; Schmidt et al., 2014). In brief, after anesthesia in tricaine, we inserted a hypodermal needle directly into the right telencephalic hemisphere while the contralateral left hemisphere was kept intact and served as a control.

For the DMH1 treatment, fish were bathed in 200 ml fresh fish water containing 10 or 20 μM DMH1 diluted from a 10 mM DMH1 (Tocris) stock solution in DMSO. For LY411575 treatment, 10 mM LY411575 (Sigma-Aldrich) stock solution was diluted with fresh fish water to 6 μM or 3 uM final concentration. For the combined treatment 200 μL of a 10 mM DMH1 stock solution (final concentration 10 μM) plus 60 μl of a 10 mM LY411575 stock solution (final concentration 3 μM) were mixed with 200 ml fresh fish water. Three fish were kept in the solutions for 5 days. Every morning, fish were fed with regular adult fish food and the fish water containing the drug was changed daily.

For heat-shock, adult zebrafish were transferred to a beaker containing fresh fish water at 33-34°C (water bath). After 30 min, the temperature was increased to 37°C and the fish were kept for 1 hour at this temperature. Afterwards, the fish were transferred to 28.5°C and kept for 6 more hours before being euthanized.

### Constructs and synthesis of antisense DIG RNA probes

The following antisense digoxigenin-labeled probes were used: *id1*, *her4.1*, *ascl1a* and *notch3* (Armant et al., 2013). The *bmp2a*, *bmp7a*, *bmp7b* (Shawi and Serluca, 2008) and *bmpr2b* (Monteiro et al., 2008) probes were amplified by PCR from zebrafish embryonic cDNA (primers see Table S3), then cloned into the pGEM-T easy vector (Promega). *bmp2b* and *bmp4* were kindly provided by Matthias Hammerschmidt (Hild et al., 2000) and *bmpr1ab* by Jeroen Bakkers (Smith et al., 2011). Briefly for probe synthesis, 1 μg of each plasmid was linearized using appropriate restriction enzymes for 30 min at 37°C. After deactivation of the restriction enzyme (see Table S4) at 80°C for 5 min, the plasmid was used for *in vitro* RNA transcription in the presence of DIG labelling mix (Roche) and RNA Polymerase (see Table S4) and incubated for 3 h at 37°C. The reaction was stopped by adding 0.2 M EDTA, pH8 and purified using the ProbeQuant G50 Micro column kit (GE Healthcare). The probe was then diluted 1:1 using hybridization buffer (Schmidt et al., 2014) for storage at −20°C.

### Preparation of adult zebrafish brains, *in situ* hybridization, immunohistochemistry, imaging and quantification

Brain preparation (dissection and sectioning) for *in situ* hybridization and immunohistochemistry were performed as described in Schmidt et al (2014).

For immunohistochemistry, primary antibodies included: chicken anti-GFP (1:1000, Aves labs, Davis, California), mouse anti-PCNA (1:500, Dako, Agilent, Santa Clara, California), rabbit anti-S100β (1:400, Dako), mouse anti-GS (Glutamin Synthetase) (1:1000, Millipore), rabbit anti-HU (1:500, Abcam), rabbit anti-BMP2b-Zebrafish (1:50, Anaspec, AS-55708), mouse anti-NeuroD1 (1:500, Abcam), rabbit anti-Phospho Smad1/5/9 (1:200, Cell Signaling Technology) and rabbit anti-RFP (1:500, antibodies online). Secondary antibodies were conjugated with Alexa fluor dyes (Alexa series) and included anti-chicken Alexa 488, anti-mouse Alexa 546 and anti-rabbit Alexa 633 (1:1000, Invitrogen, Waltham, Massachusetts). For in situ hybridization, the prepared DIG probes were hybridized with the brain tissue. After cutting, secondary DIG antibodies (anti-DIG-AP for chromogenic staining; anti-DIG-POD for fluorescent staining) were applied overnight. Staining took place on the next day with NBT/BCIP in the case of chromogenic staining or Tyramide Cy3 solution (Perkin Elmer) for fluorescent staining. Pictures of chromogenic *in situ* hybridized sections were acquired with a Leica stereomicroscope MZ16 F. Immunohistochemically stained brain slices were mounted using Aqua-Poly/Mount (Cat No. 18606-20, Polysciences, Inc) with coverslips (0.17mm thickness) and imaged with a laser scanning confocal microscope (Leica TCS SP5). To obtain single-cell resolution images, an HCX PL APO CS x63/1.2NA objective was used with the pinhole size set to 1-airy unit. Fluorescent images for green (GFP), red (PCNA), and infrared channesl (S100β, GS, HU, Bmp2b, NeuroD1, pSmad1/5/9, and mCherry) were acquired sequentially in 16-bit color depth with excitation/emission wavelength combinations of 488◻nm/492-550◻nm, 561◻nm/565-605◻nm, and 633◻nm/650-740◻nm, respectively. Pixel resolution for XY and Z planes are 0.24 and 0.50◻μm, respectively. For individual brain samples, at least three transverse sections cut with a vibratome (VT1000S, Leica) at different anterior-posterior levels representing anterior, posterior an intermediate telencephalic regions were imaged.

### Quantitative real-time PCR

Total RNA was isolated from adult telencephala using Trizol (Life Technology). First-strand cDNA was synthesized from 1 μg of total RNA with the Maxima First-Strand cDNA synthesis kit (Thermo Scientific) according to the manufacturer’s protocol. A StepOnePlus Real-Time qRT-PCR system (Applied Biosystems) and SYBR Green fluorescent dye (Promega) were used. Expression levels were normalized using β-actin. The relative levels of mRNAs were calculated using the 2^−ΔΔCT^ method. The primer sequences (Mitra et al., 2019) are listed in Table S5. Experiments were performed at least 3 times, each time with RNA pooled from 5 brains for WT or *id1*^*ka706/ka706*^, respectively.

### Statistical analysis

For quantifications, the number of cells was determined by counting the cells in Z-stacks of 50◻μm thickness in 1 μm steps (×63 objective). Three sections per brain were analyzed.

For quantification of the changes in *id1* and *her4.1* mRNA levels following chemical treatment, as well as *ascl1a*, pictures were taken with a stereo microscope and then processed by ImageJ to set up a threshold for the staining intensity. Quantification of the staining intensity was performed for control and chemical treatment from the dorsomedial to the dorsolateral part of the telencephalon. Three brains were analyzed, and the fold-induction between the experimental group vs control was calculated.

Comparisons between two data sets and between more than two data sets were performed using Welch two-sample t-test and one-way ANOVA followed by Tukey’s multiple pairwise-comparison test, respectively. Statistical significance was assessed using R.

### Image analysis

Confocal brain images were opened with Fiji/ImageJ software as composite hyperstacks to manually evaluate colocalization of GFP, PCNA, S100β, GS, HU, mCherry proteins and expression of *bmp* genes and *notch3* mRNA (FISH). Cells expressing individual markers or marker combinations were counted in the dorsomedial and the dorsolateral ventricular zones in three transverse sections prepared at different anteroposterior levels of the telencephalon.

### Generation of the *id1* knockout allele *id1*^*ka706*^

The oligonucleotide containing the target sequence (5’-CCAAAATGAAAGTTGTGGGACCT-3’), was designed using the ZiFiT Targeter program (Hwang et al., 2013). The guide RNA was synthesized using the cloning-free guide RNA synthesis method adapted from (Gagnon et al., 2014) where an oligonucletide containing the T7 promoter is annealed to the gene specific target oligo. After annealing, T4 DNA Ligase was added and the mixture was incubated at 12°C for 20 min in a thermocycler to fill up the sequence of the annealed oligonucleotides to a double stranded DNA. Afterwards, the DNA was purified by column purification (Gel and PCR clean up kit, Macherey-Nagel) and used for RNA synthesis (Megashortscript T7 Kit, Ambion).

Single-cell stage embryos were injected with 300 ng/μl Cas9 protein (GeneArtTM PlatinumTM Cas9 Nuclease; Invitrogen/Life Technologies) along with 200 ng/μl of previously synthesized guide RNA and phenol red to a final concentration of 0.05% as visual marker for injection. F0 embryos were raised to adulthood and then outcrossed with WT animals. F1 progeny with indel mutation were in-crossed, and homozygous F2 mutants were identified.

For genotyping, genomic DNA was isolated from injected embryos or fin biopsies from adult fish by the HotSHOT method (Meeker et al., 2007) for determination of guide RNA efficiency as described in (Etard et al., 2017). Genomic DNA was prepared by incubating biopsy samples in 75 μl of 25 mM KOH with 0.2 mM EDTA at 96°C for 20 min, followed by neutralisation with 75 μl of 40 mM Tris-HCl (pH 3.8). The genomic region containing the site of mutation was PCR-amplified using the following conditions: initial denaturation step at 94°C for 7 min; 35 cycles of 94°C for 25 s, 56°C for 30 s and 72°C for 30 s; and a final elongation step at 72°C for 10 min. For *id1*, a 451 bp amplicon encompassing the mutation site was generated using the following primers: forward 5’-CATCATCCGCAGAAGACACA-3’; reverse 5’-AACATGGTCATCTGCTCGTC-3’. The PCR product was sent for Sanger sequencing to identify the mutant alleles (Microsynth).

### Single-cell sequence analysis

For assessing genes co-expressed with *id1* in radial glia cells, we downloaded the prepared count data of 370 cells (zebrafish_neurogenesis_smartseq.h5ad in https://github.com/fabianrost84/lange_single-cell_2019). We used the Scanpy package to read this file (Wolf et al., 2018). After the quality control to remove the low-quality cells, 264 cells were used for further analysis. The number of radial glia cells was 76. The number of radial glia cells with *id1* expression was 44. The set of 44 *id1+* RGC cells contained 22 quiescent (ccnd1-negative) cells. We calculated the Pearson’s correlation coefficient between *id1* gene expression and expression of other genes in *id1* and *ccnd1*-RGCs by R function (cor.test).

## Supplementary figures

**Figure S1. BMP2b is not detectably expressed in oligodendrocytes of the telencephalon.** Cross-sections through the telencephalon of *Tg(olig2:gfp)* transgenic fish marking oligodendrocytes (green) were immunostained with a BMP2b antibody (blue). The boxed-in area in A is magnified in A’ to A’’’ showing individual channels (A’, A’’) and the merged image (A’’’). White arrows show 2 oligodendrocytes, yellow arrows show BMP2b+ neurons. Scale bars: 100 μm (A), 20 μm (A’-A’’’).

**Figure S2. Heat-shock does not influence the expression pattern of *id1* in WT fish.** (A-C) Cross-sections through telencephala of heat-shocked WT, (A) and *hs:dnBMPR1a* transgenic animals without (B) and with heat-shock (C). Heat-shock of WT animals did not influence the expression of *id1*. In contrast, decrease of *id1* expression was noted after inhibition of the BMP pathway by heat-shock of *Tg(hs: dnBmpr1a)* animals (C) relative to transgenics without heat-shock (B). Black arrows show the expression of *id1* in the ventricular zone of WT (A) and *Tg(hs:dnBmpr1a)* animals without heat-shock (B) and after heat-shock (C). n=3 animals (A-C). Scale bar: 20 μm. HS, heat-shock

**Figure S3. Both *notch3* and *her4.1* are expressed together with *id1* in the RGCs of the telencephalon.** (A-A’’) Fluorescent ISH against *notch3* mRNA (A, red) on transverse sections through the telencephalon of *Tg(id1-CRM2:gfp)* transgenic animals (A’, green). A’’ merged view of panels A and A’. An overlapping pattern of expression for *notch3* and *Tg(id1-CRM2:gfp)* (green) was noted in the telencephalon (white arrows). (B-C’’’) Immunostaining on cross sections of the telencephalon of *Tg(her4.1:gfp;id1-CRM2:mCherry)* double transgenics with antibodies against GFP (green), mCherry (blue) and the proliferation marker PCNA (red). GFP and mCherry signals co-localize indicating that her4.1 and id1 are co-expressed. B’’’ inset: magnified image of a PCNA+/id1-CRM2 : mCherry+ cell (yellow arrowhead, upper image) and a PCNA+/her4.1+ cell (white arrowhead, lower image). C’’’ inset: a magnified view of a PCNA+/id1-CRM2:mCherry+/her4.1:gfp+ cell (red arrowhead). D: Summary of co-expression analysis of (*Tg(id1-CRM2:mCherry)*, red) and (*Tg(her4.1:gfp*), green). Pie charts represent total number of *Tg(her4.1:gfp)* or *Tg(id1-CRM2:mCherry)* or both *Tg(her4.1:gfp)* and *Tg(id1-CRM2:mCherry)* expressing cells and the fraction of cells co-expressing PCNA. Cells expressing either *Tg(id1-CRM2:mCherry)* alone or *Tg(her4,1:gfp)* alone show a higher proportion of PCNA+ cells than cells expressing both markers. Scale bar = 20 μm. n=3 brains.

**Figure S4. Expression of *id1* is independent of Notch signaling** (A-C) Expression of *id1* revealed by ISH on control brains (DMSO, A) or brains treated with different concentrations of DMH1 (B) or LY (C). (B) Inhibition of BMP signaling by 20 μM DMH1 lead to the reduction of *id1* expression. (C) Inhibition of Notch signaling by 6 μM LY411575 did not influence the expression of *id1* but strongly reduced *her4.1* expression (See Fig, 4E) Black arrows show the expression of *id1* in the ventricular zone of untreated (A) and treated (B and C) adult zebrafish telencephala. n=3 brains. Scale bar: 100 μm.

**Figure S5. Inhibition of BMP signaling induced reduction of *her4.1* expression.**

(A-B) ISH reveals decrease of *her4.1* expression after inhibition of the BMP pathway by heat-shock of *Tg(hs:dnBmpr1a)* animals. Black arrows show the expression ofthe *her4.1* gene in the ventricular zone of *Tg(hs:dnBmpr1a)* animals without (A) and with heat-shock (B). (C) Quantification of *her4.1* expression in panels A and B. n=3 independent brains (A-B), n=15 sections (C). HS, heat-shock

**Figure S6. Generation of a CRISPR/Cas 9-mediated Id1 loss of function mutant *id1***^***ka706/ka706***^. (A) Schematic representation of the *id1* locus on chromosome 11. The gene consists of 2 exons. Red letters represent the start codon of the *id1* coding sequence. The Sanger sequencing results of WT (*id1*^*WT*^) and homozygous (*id1*^*ka706/ka706*^) sequences are displayed below. In the homozygous sequences, T and G of the start codon are deleted. (B) Quantification of the S100β+ RGCs in WT and *id1*^*ka706/ka706*^ telencephala without injury. n=3 brains (B).

**Table S1.**
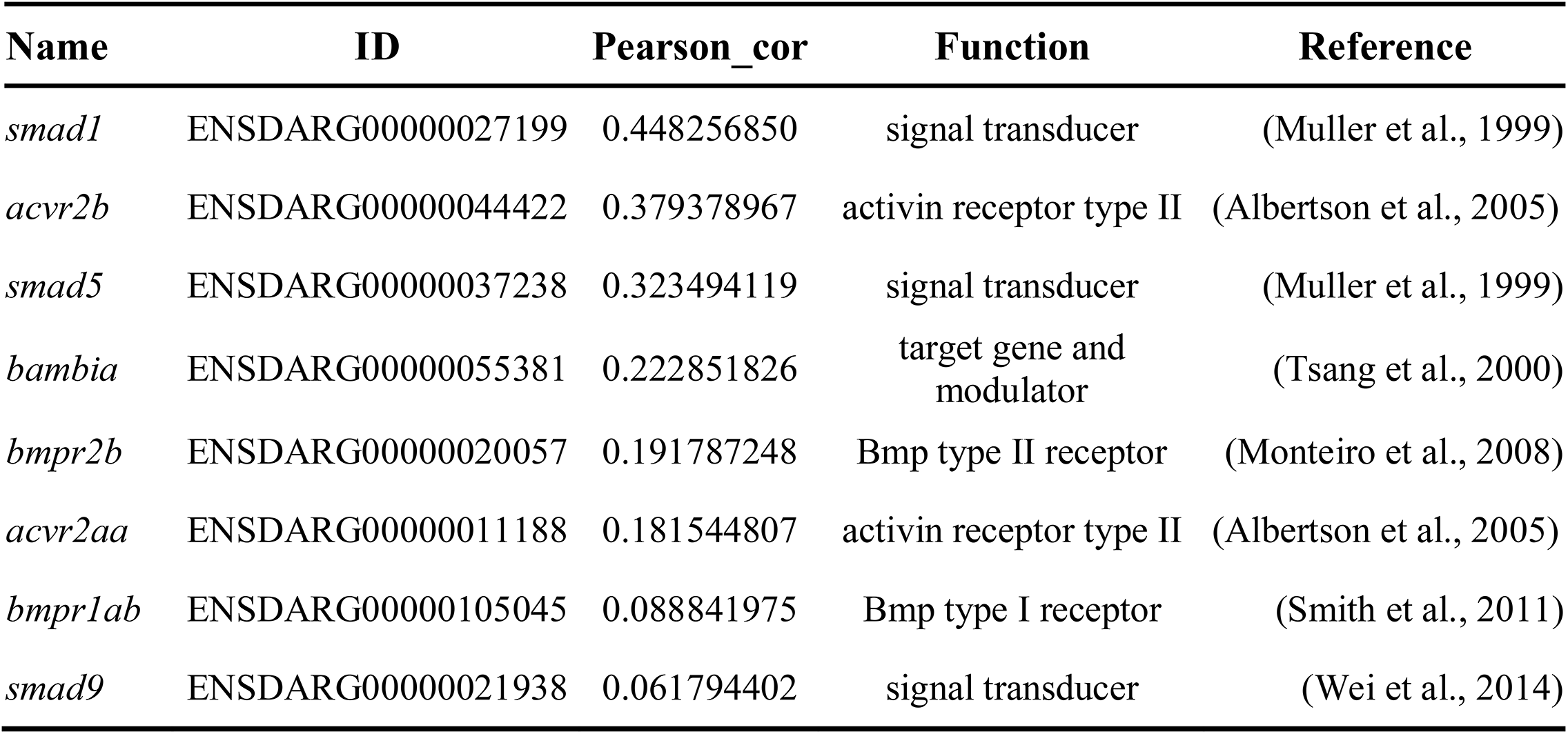
Genes co-expressed with *id1+*/*ccdn1*- RGCs according to single cell sequencing data from Lange et al., (2020).

**Table S2.**
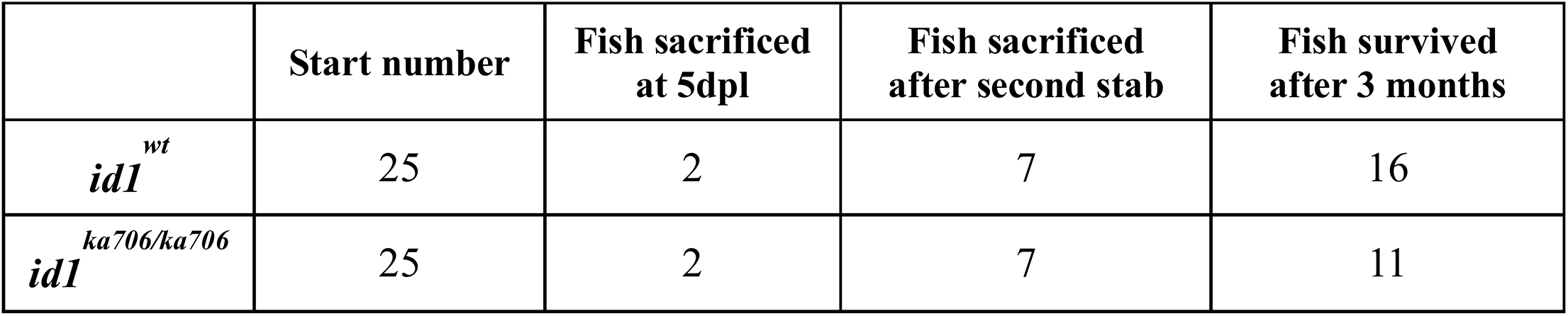
Number of WT and *id1*^*ka706/ka706*^ adult zebrafish used in the multiple stab wound experiments.

**Table S3.**
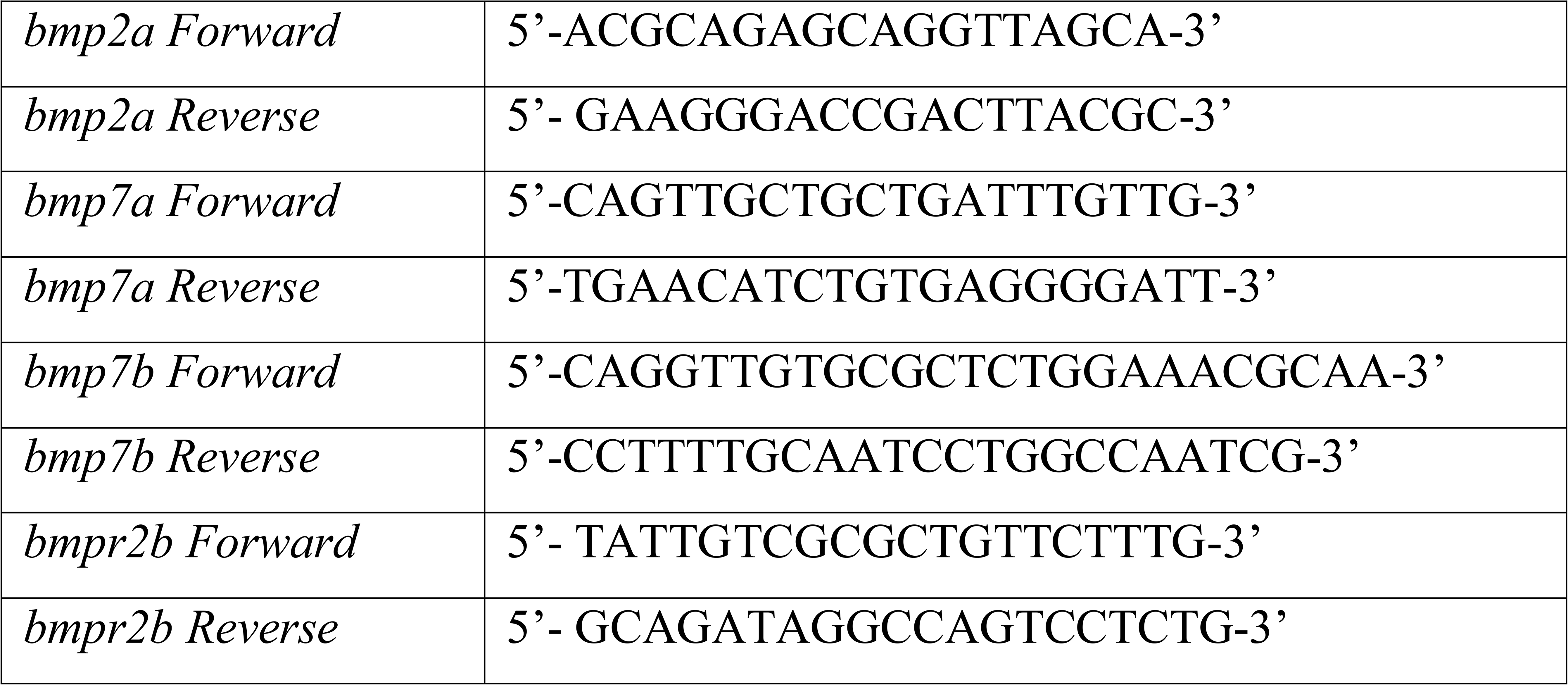
PCR primer sequences for ISH probes.

**Table S4.**
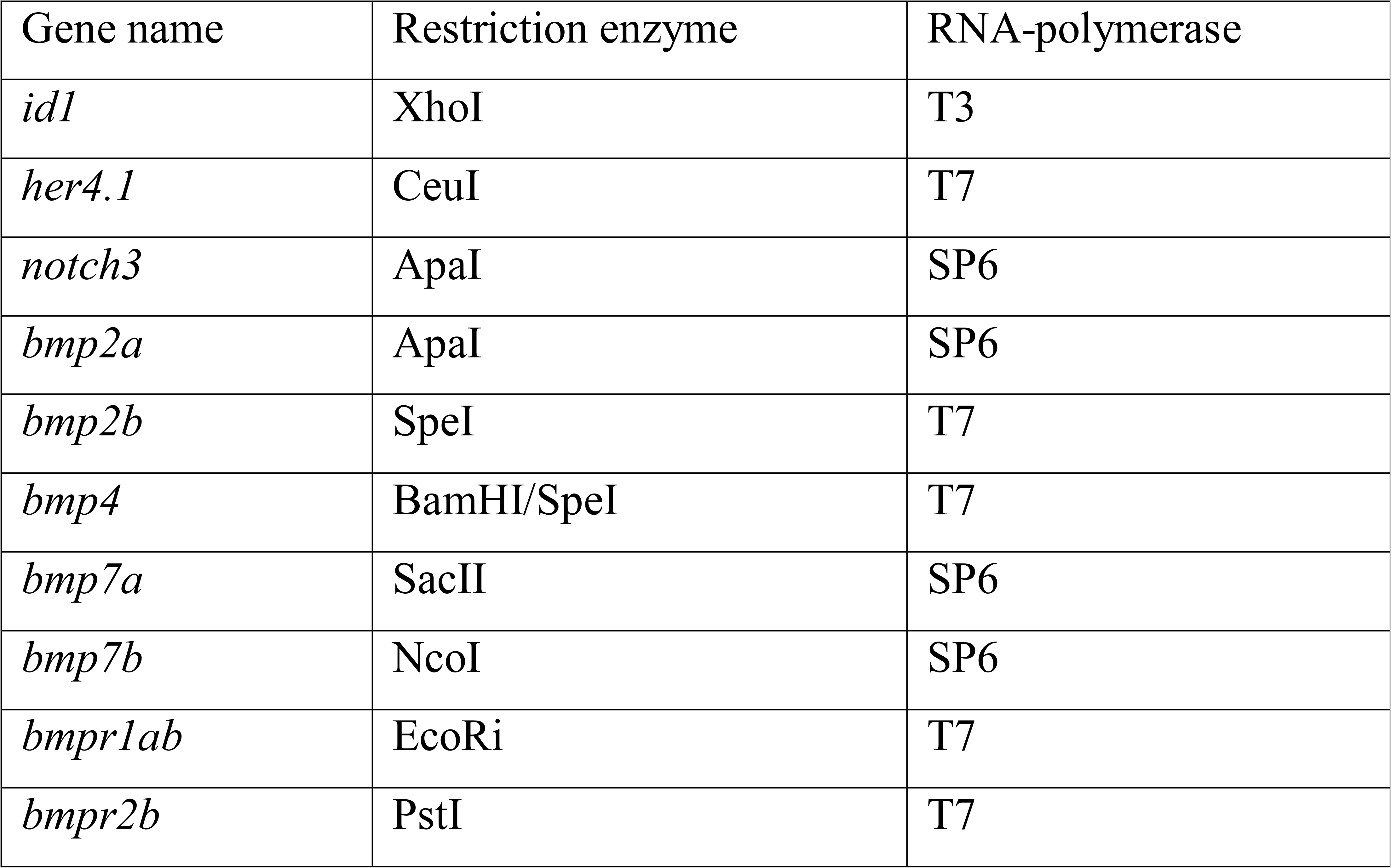
Restriction enzymes and RNA polymerases used for DIG probe synthesis.

**Table S5.**
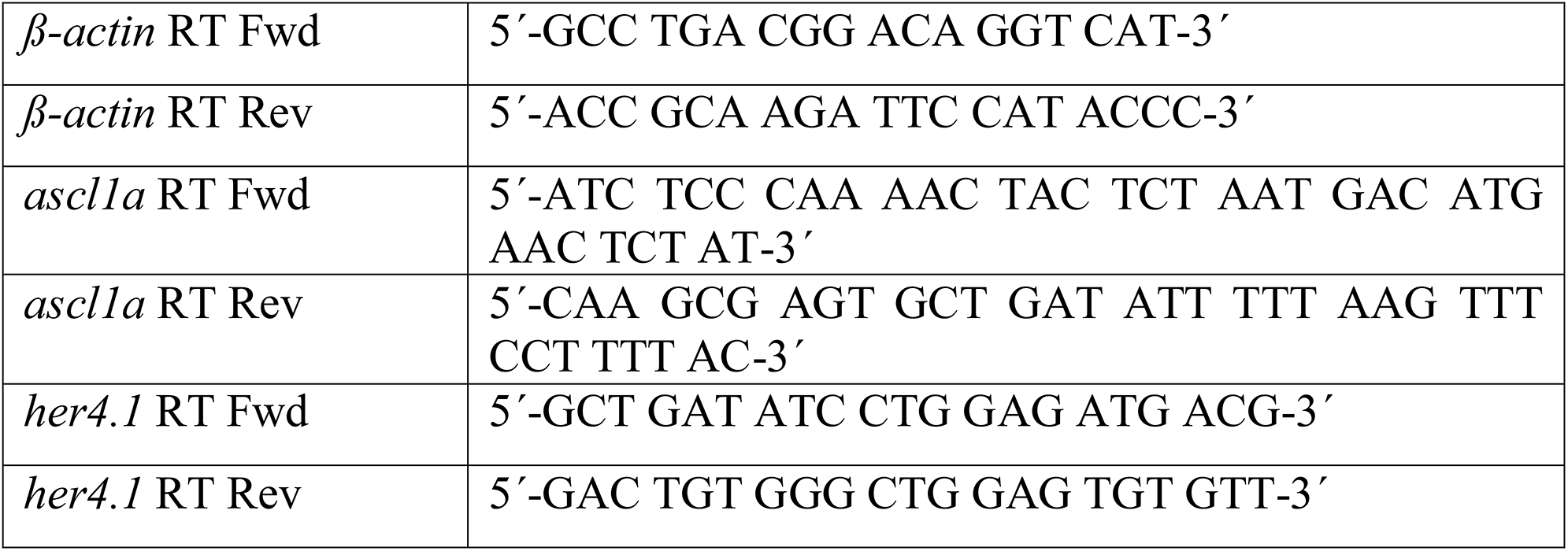
PCR primer sequences for RT-qPCR.

